# Sex-biased effect of sodium leak channel NALCN deletion in striatal *Drd2* spiny projection neurons

**DOI:** 10.1101/2024.10.23.619832

**Authors:** Laia Castell, Claire Naon, Angelina Rogliardo, Leila Makrini, Maëlle Avrillon, Audrey Mignon, Claire Bernat, Philippe Lory, Federica Bertaso, Arnaud Monteil, Clémentine Bosch-Bouju, Emmanuel Valjent

**Affiliations:** IGF, Univ. Montpellier, CNRS, Inserm, F-34094 Montpellier, France; Department of Physiology, Faculty of Medicine Siriraj Hospital, Mahidol University, Bangkok, Thailand; Université Bordeaux, INRAE, Bordeaux INP, NutriNeuro, UMR 1286, Bordeaux, France

**Keywords:** sodium leak channel, striatum, D2R spiny projection neurons, intracellular signalling, motivated-behaviors

## Abstract

The sodium leak channel NALCN is an important modulator of neuronal excitability, yet its specific role in striatal medium-sized spiny neurons remains largely unexplored. In this study, considering that *Nalcn* transcripts are enriched in the dorsal and ventral striatum of *Drd2*-SPNs, we investigated the functional impact of NALCN deletion in *Drd2*-expressing SPNs in both male and female mice. Electrophysiological recordings revealed significant sex differences, with male SPNs exhibiting altered membrane properties and increased excitability, while females showed more subtle changes. Interestingly, eticlopride-induced intracellular signaling was selectively enhanced in female SPNs lacking NALCN. Behaviorally, male mice exhibited reduced motivation in food-seeking tasks and impaired discrimination of threat cues. Our findings uncover an important, sex-specific role for NALCN in regulating striatal function and behavior and underscore its significance in maintaining normal striatal function.

## INTRODUCTION

The dorsal striatum (DS) and its ventral part, the Nucleus Accumbens (Acb), are subcortical structures involved in adaptive control of behavior (Balleine et al., 2009; Klawonn & Malenka, 2018). While the DS is essential for motor control, learning of habits and skills, the NAc processes incentive-reward responses associated with novel, hedonic, stressful or aversive stimuli (Floresco, 2015; Yin & Knowlton, 2006). Through the fine adjustment of a filtering process, the GABAergic medium-sized spiny neurons (SPNs) select relevant sensory cues predicting the occurrence of positive or negative events to favor the optimization of action plans maximizing therefore reward outcomes (Robbe, 2018; Yin & Knowlton, 2006). Consequently, imbalanced activity of SPNs has been associated with profound striatal dysfunctions causally linked to some of symptoms found in various neurological, psychiatric and neurodevelopmental disorders (Bateup et al., 2010; Kravitz et al., 2010).

Despite a powerful synaptic excitation driven by cortical and thalamic inputs, SPNs display a low rate of firing activity resulting from a hyperpolarized resting membrane potential (Wilson & Kawaguchi, 1996). This unique integrative profile of SPNs is mainly due to the expression of voltage-dependent potassium channels generating inward rectifying currents (Nisenbaum & Wilson, 1995; Uchimura et al., 1989). All SPNs, regardless they express dopamine D1 (*Drd1*-SPNs) or D2 (*Drd2*-SPNs) receptors, in both male and females, share this electrophysiological feature (Godino et al., 2023; Moyer et al., 2007). However, differences in excitability between SPNs sub-populations have been observed, depending on the developmental stage, the sub-territory, and experience-driven plasticity events (Cao et al., 2018; Deroche et al., 2020; Planert et al., 2013). These variations most likely result from the different expression levels of ion channels repertoire recently unveiled by high-throughput RNAseq (Montalban et al., 2022; Puighermanal et al., 2020), and the complex neuromodulation occurring in the striatum.

The sodium leak channel NALCN is widely expressed throughout the brain. It regulates the resting membrane potential of neurons and subsequently their excitability (Monteil et al., 2024). Its activity is finely regulated both positively and negatively by neuromodulators through G protein-coupled receptors and its dysfunction is associated with disease states in human (Cochet-Bissuel et al., 2014; Monteil et al., 2024). Enriched in neuronal cell types displaying tonic firing, NALCN conductance has been shown to be essential to sustain the pacemaker activity of GABAergic neurons of the *substantia nigra pars reticulata* (Delgado-Zabalza et al., 2023; Lutas et al., 2016) and midbrain dopamine (DA) neurons (Cobb-Lewis et al., 2023; Philippart & Khaliq, 2018; Um et al., 2021). However, although highly expressed in SPNs, predominantly in *Drd2*-SPNs, a comprehensive understanding of the role of NALCN-mediated currents in these highly hyperpolarized neurons with low firing rate is still lacking.

In this study, we investigated the functional consequences of *Nalcn* invalidation in striatal *Drd2*-SPNs in both male and female mice. We unveiled sexual dimorphic effects *i*) on passive membrane properties and excitability of *Drd2*-SPNs, *ii*) on the regulation of signaling events occurring in *Drd2*-SPNs in response to a single administration of the D2R antagonist eticlopride and *iii*) on the regulation of striatal-dependent behavior such as food-seeking or dynamic defensive behaviors. Together, our results uncover previously unknown function of sodium leak channel NALCN in striatal D2R neurons.

## METHODS

### Animals

Male and female C57Bl/6J (Charles River Laboratories, France) were used for *in situ* hybridization. Male and female resulting from breeding *Drd2^Cre/+^* with *Ai95^f/f^* mice were used as control for immunofluorescence analysis and patch-clamp recording. The deletion of *Nalcn* from D2R neurons was obtained by crossing *Drd2^Cre/+^* mice with *Nalcn^f/f^* (Flourakis et al., 2015) (kindly provided by Pr D Ren) or *Ai95 ^f/f^;Nalcn^f/f^* mice. Male and female *Drd2^Cre/+^;Ai95 ^f/f^;Nalcn^f/f^* mice (cKO) were used for immunofluorescence analysis and patch-clamp recording. Behavioral experiments have been performed on *Nalcn^f/f^* (Cre-negative, control) and *Drd2^Cre/+^;Nalcn^f/f^*(Cre-positive, cKO) male and female mice. All the behavioral experiments have been performed on independent cohorts of mice. All mice were housed in groups of 2 to 5 per cage (standard sizes according to the European animal welfare guidelines 2010/63/EU) and maintained in a 12h light/dark cycle (lights on from 7:00 am to 7:00 pm), in stable conditions of temperature (22°C) and humidity (60%), with food and water provided *ad libitum*. All animal procedures were conducted in accordance with the guidelines of the French Agriculture and Forestry Ministry for handling animals (authorization number/license B34-172-41) and approved by the relevant local and national ethics committees (authorizations APAFIS#22635).

### Single molecule *in situ* hybridization

Expression of *Nalcn*, *Drd2*, *Drd1* transcripts were detected using single molecule fluorescent *in situ* hybridization (smFISH) (Castell et al., 2024). Brains from 3 C57BL/6 male and female mice were rapidly extracted and snap-frozen on dry ice. Coronal sections (16 µm) of the striatum comprising the dorsal striatum (SD) and nucleus accumbens (Acb) (bregma 1.34 to 1.10 mm) were collected directly onto Superfrost Plus slides (Fisher Scientific). Probes for *Drd2* (ACDBio; Mm-drd2-C3, Cat# 406501-C3), *Drd1* (ACDBio; Mm-drd1-C2, Cat# 406491-C2) and *Nalcn* (ACDBio; Mm-nalcn, Cat#415161) were used with the RNAscope Fluorescent Multiplex Kit (ACDBio; Cat# 323110). After incubation with fluorescent probes, slides were counterstained with DAPI. ProLong Diamond Antifade mountant was used as mounting medium (Thermo Fisher Scientific catalog #P36961). Fluorescent images of labeled cells were captured using sequential laser scanning confocal microscopy (Leica SP8) at the Montpellier RIO imaging facility.

### Immunofluorescence

Tissue preparation and immunofluorescence analyses were performed as previously described (Cutando et al., 2022). Mice were rapidly anaesthetized with Euthasol® (360 mg/kg, i.p., TVM lab, France) and transcardially perfused with 4% (weight/vol) paraformaldehyde in 0.1 M sodium phosphate buffer (pH 7.5). Brains were post-fixed overnight in the same solution at 4°C. Coronal striatal sections (40-µm thick sections) were obtained using a vibratome (Leica, France). Slices were stored at -20°C in a solution containing 30% (vol/vol) ethylene glycol, 30% (vol/vol) glycerol and 0.1 M sodium phosphate buffer (PBS), until they were processed for immunofluorescence. On day 1, selected free-floating sections were rinsed three times in PBS then incubated 15 min in 0.2% Triton X-100 in PBS and blocked for 1 hour in 3% bovine serum albumin (BSA) in PBS before incubation 72 hours at 4°C with the primary antibodies (**Supplemental Table 1**). On day 2, slices were rinsed three times in PBS and incubated 45 min with goat Cy3-coupled anti-rabbit (1:500; Jackson ImmunoResearch Laboratories), goat Alexa Fluor 488-coupled anti-mouse or chicken and goat Alexa Fluor 647-coupled anti-rat (1:500; Jackson ImmunoResearch Laboratories) secondary antibodies. Sections were rinsed twice in TBS and twice in 0.25 M Tris-buffer before mounting in Surgipath Micromount (Leica). Single and double-immunolabeled images were single confocal sections captured using sequential laser scanning confocal microscopy (Leica SP8) at the Montpellier RIO imaging facility.

### Western Blot

Mice were sacrificed by cervical dislocation, their heads were cooled in liquid nitrogen for 4□s, and the brains were removed (Cutando et al., 2022). The striata were dissected out on an ice-cold surface, sonicated in 300□μl of 10% SDS, and boiled at 100°C for 10□min. Aliquots (5□μl) of the homogenate were used for protein determination using a BCA assay kit (Pierce) (Lot# RG235624; Thermo Fisher Scientific). Equal amounts of striatal lysates (20□μg) for each sample were separated by 11% SDS-polyacrylamide gels before electrophoretic transfer onto Immobilon-P membranes (Millipore, (#IPVH00010). Membranes were blocked for 30 min with 4% bovine serum albumin (BSA) in 0.1 m PBS before incubation for 2 hours with primary antibodies (**Supplemental Table 1**). Detection was performed using horseradish peroxidase-conjugated antibodies to rabbit or mouse (1:10,000; Cell Signaling Technology) and visualized by enhanced chemiluminescence detection (Luminata Forte Western HRP Substrate; Millipore, # WBWF0500). The optical density of the relevant immunoreactive bands was quantified after acquisition on a ChemiDoc XRS System (Bio-Rad) controlled by Image Lab software version 3.0 (Bio-Rad). Signals were normalized to β-actin or GAPDH values in the same sample and expressed as a percentage of control treatment or group.

### Behaviors

#### Locomotor activity

Horizontal and vertical activity was measured in a circular corridor (Imetronic, Pessac, France) for 90 min. Horizontal locomotor activity was counted as travels through one-fourth of the circular corridor as detected by consecutive interruption of two adjacent beams (1/4 turns). Vertical activity represented the number of rearings detected by the interruption of beams placed at a height of 7.5 cm along the corridor

#### Beam walking test

Fine motor coordination was assessed by using the beam-walking test performed as previously described (Cutando et al., 2022). Mice were trained two consecutive days to cross a horizontal wooden circle rod (1□m long, 3□cm in diameter) placed 60□cm above a cushioned table. For the test day, mice crossed twice a thick wooden rod (3□cm in diameter) and a thin wooden rod (1□cm in diameter). The time spent in the beams and the total number of footslips were recorded for each mouse.

#### Elevated plus maze

Elevated plus maze was performed as previously described (Puighermanal et al., 2020). The number of entries and time spent in the closed arms (4 paws within closed arm) or open arms (4 paws within open arms) were calculated during 5 min.

#### Marble burying and Digging test

Marble burying and digging test were performed as previously described (Puighermanal et al., 2020). Marble burying test was conducted two consecutive days. For the digging test, parameters evaluated included the duration of digging, the number of digging bouts and the latency to initiate the first digging bout.

#### Food-seeking behavior

The experiment was carried out using soundproof polymodal system (Imetronic, Pessac, France) as previously described (Castell et al., 2024). A week before the beginning of the conditioning phase, animals were individually housed and food-restricted to maintain 85% of their original body weight. Food restriction was maintained during the integrality of the experiment, which was performed during the light phase of the dark/light cycle. High palatable isocaloric pellets of 20 mg (TestDiet), similar in caloric content to the standard diet (3.48 kcal/g) but with higher level of sucrose among the carbohydrates (49%) and with the addition of chocolate flavor were used as reinforce. The experiment started with a fixed ratio (FR)-1 schedule of reinforcement from day 1 to day 4. Mice had to lever press once in the active lever to receive a pellet. Each lever press leading to a pellet reward in the active lever was followed by a 15-second time-out period in all the phases of the experiment. FR1 schedule was then followed by 3 days under FR5 (day 5-7), where 5 consecutive lever presses were needed to obtain one pellet. On day 8, mice underwent a progressive ratio (PR) schedule of reinforcement in which the number of active lever presses required before reinforcer delivery exponentially increased.

#### Discriminative active avoidance

The experiment was performed in a soundproof shuttle box (Imetronic, Pessac, France) as previously described (Castell et al., 2024). On day 1, mice underwent a habituation session consisting of 10 presentations of two different tones, a CS- and CS+ lasting 13 sec maximum. Mice can stop both CSs by shuttling from one compartment to another. From day 2-4 (session 1-3), mice were underwent a training session (once per day) during which they were exposed to CS+/CS-in a pseudo-random manner. Again, the CS (+ or -) stops whenever the mice crosses from one compartment to another. CS-last 8 sec maximum. CS+ last 13 sec maximum, but after 8 sec the unconditioned stimulus (0.7 mA foot-shock coinciding with the last 5 sec of the CS+ presentation) is delivered until the mice shuttles or for a maximum of 5 sec. During the inter-trial period, the animal was free to cross between compartments. The probabilities of avoidances during CS+ and defensive responses (escape + avoidance) during CS-were used as an index of learning and discrimination, respectively.

#### Discriminative auditory fear conditioning

The experiment was carried out in soundproof fear conditioning apparatus (Imetronic, Pessac, France) as previously described (Castell et al., 2024). On day 1, mice underwent a period of habituation to box A or B during which two different tones (CS+ and CS-; 30 sec 4 times each) were randomly presented. The conditioning session occurred later the same day. Mice were placed on the same box as for the habituation and after 5 min of exploration they received 5 pairings of the CS+ (30 sec) with the foot-shock at the last sec of the CS+ (1 sec, 0.6 mA), and 5 presentations of the CS-alone (30 sec). Interval between each tone was a random time between 20 and 180 sec. In the habituation and conditioning phases, cages were cleaned with 70% ethanol. On day 2, mice were tested in a box different from the one used for conditioning. During this test session, mice received 12 presentations of the CS+ (30 sec) and 4 presentations of the CS-. The inter-trial interval in all sessions was between 20 and 180 sec. During this session, boxes were cleaned with 1% acetic acid. The meaning of each tone (pairing or not with the foot-shock) was counterbalanced between mice and genotypes. Freezing behavior was recorded using a tight infrared frame. The threshold for considering freezing behavior was set up at 2 sec. The first 10 min of habituation were used to assess the basal freezing. Freezing probability was calculated during CS+ and CS-presentation and were averaged by blocks of 4 CS (+ or -) presentation.

### Patch-clamp electrophysiology

#### Slice preparation for patch-clamp recordings

Striatal brain slices were prepared as previously described (Bosch-Bouju et al., 2016). Briefly, following deep anaesthesia with isoflurane, brain was quickly extracted from the skull and plunge in artificial cerebrospinal fluid (ACSF) containing (in mM): 125 NaCl, 25 NaHCO3, 2.5 KCl, 1.25 NaH2PO4, 2 CaCl2, 1 MgCl2, 25 glucose and 1.25 pyruvate, bubbled with carbogen gas. Sagittal slices (300 µm) were obtained with a vibratome (VT1000S, Leica) in cold and oxygenated ACSF solution. After slicing, slices were immediately transferred in ACSF solution at 34°C for 1h, then at room temperature for at least 1 hour before recording.

#### Neurons identification and patch-clamp recordings

Slices were recorded at 30°C while continuously superfused at 1.5-2 ml/min with oxygenated ACSF (peristaltic pump minipulse 3, Gilson). Neurons were visualised under an upright fluorescent microscope (Nikon FN1) with 40X water-immersion objective. Fluorescence (GCaMP6) was detected with pE-300 and CCD camera (INFINITY 3S-1UR M/C). For each recorded neuron, fluorescence was detected during the first protocol eliciting spikes. A neuron with detectable fluorescence at the moment of spikes was categorized as ‘*Drd2*-positive neuron’ while neurons with no detectable fluorescence was categorized as ‘*Drd2*-negative neuron’. In every case, we confirmed that fluorescent neurons were detectable in the area of the patched neuron. The electrophysiological profile was used to determine whether the recorded neuron was an SPN or not. Only SPNs were considered in this study. Patch pipettes (5-6 MΩ) were pulled from borosilicate glass capillaries (GBF-150-117-10; Sutter Instruments) with a micropipette horizontal puller (P-97, Sutter Instruments). Electrophysiological recordings were performed using a MultiClamp 700B amplifier (Molecular Devices) and acquired using a Digidata 1550B digitizer (Molecular Devices), sampled at 20 kHz for current clamp and 200 kHz for voltage-clamp, and filtered at 1-2 kHz. All data acquisitions were performed using pCLAMP 11.0.3 software (Molecular Devices). Stimulation of excitatory fibers was performed with a stimulating electrode (bipolar concentric electrode from Phymep and stimulator A365, World Precision Instruments) located in the fiber bundle between the cortex and the ventral striatum. Intracellular solution based on K-gluconate (containing in mM: 128 KGlu, 20 NaCl, 1 MgCl2, 1 EGTA, 0.3 CaCl2, 2 Na2-ATP, 0.3 Na-GTP, 0.2 cAMP, 10 HEPES, pH 7.35) was used for all recording protocols.

#### Recording protocols and analysis

Neurons were monitored in both voltage-clamp and current-clamp modes. Protocols performed in voltage-clamp were: spontaneous excitatory post-synaptic currents (sEPSC), paired-pulse stimulations and current steps. Protocols performed in current-clamp were: electrophysiological profile and excitatory post-synaptic potential (EPSP)/spike coupling. Data were analysed offline using Clampfit 11.0.3 (Molecular Devices).

#### Electrophysiological profile

Recording of current/voltage curves was performed in current clamp mode at resting membrane potential, with a series of 600 ms-duration depolarizing current steps starting at -150 pA with 10 pA increment. The resting membrane potential was measured as the average potential before the step of current. Resistance was measured with the -100 pA step. Rheobase was defined as the first step with one or more spikes.

#### Voltage steps

Neurons were clamped at -70 mV without any drug in the extra or intracellular medium. Steps of voltage starting at +80 mV with an increment of -10 mV were applied for 1 sec at 0.5 Hz. Based on (Chua et al., 2020), we analysed the instantaneous and the steady state current at each step. Both currents were plotted against applied voltage and the conductance was measured as the slope of the linear regression.

#### sEPSC recordings

Neurons were clamped at -70mV in voltage clamp mode and recorded for 1 min. Events were analysed for the total duration of the protocol, except when too instable. For analysis, signal was low-pass filtered at 600 Hz (Gaussian) and electrical interference were removed (50 Hz). The event detection was made with template search function, with a template match threshold of 4, no fitting and taus calculated from 50% of peak. Event detection was verified and corrected manually. Data extracted from analysis were: sEPSCs amplitude, mean number of events/s, inter-event interval, rise time and decay time.

#### Paired-pulse stimulation

Neurons were clamped at -70 mV. Excitatory fibres were stimulated twice with an interval of 50 msec, while EPSC were monitored in voltage clamp at 0.1 Hz. A step of -5mV was applied at each sweep to monitor resistance; a variation of more than 20 % during the protocol lead to the rejection of the experiment. The step of voltage was also used to measure the capacitance of neurons. Thirty sweeps were averaged for each neuron. Amplitude of each peak was measured and paired-pulse ratio (PPR) was measured as peak amplitude 2/peak amplitude 1.

#### EPSP/Spike (E-S) coupling

Neurons were current clamped at -60 mV and stimulations of excitatory inputs were applied at 0.125 Hz with 6 different intensities (5 sweeps / intensity) chosen by the experimenter. EPSP slopes were measured off-line and the firing probability was plotted as a function of the EPSP slope (mV/ms).

### Statistics

Statistical analyses were performed with GraphPad Prism v7.0. Data were compared between *Drd2^Cre/+^;Ai95^f/+^;Nalcn^f/f^* and *Drd2^Cre/+^;Ai95^f/+^* groups, male and female groups being analyzed independently. Normality of data was tested for each set of data. Significance threshold was set at p<0.05. For variables measured with the electrophysiological profile, sEPSC and the paired-pulse ratio, statistical difference between groups was tested using Student’s t test (unpaired, two-tailed), or the non-parametric equivalent Mann-Whitney test when appropriate. Electrophysiological data represented as curves (instantaneous and steady-state currents, I/V curves, I/Spikes curves, cumulative distribution of sEPSCs amplitude and inter-event intervals) were analyzed with two-way repeated measure ANOVA. For E-S coupling, comparison of survival curves was performed with Log-rank (Mantel-Cox) test. Immunofluorescence were analyzed with two-way ANOVA. Behaviors were analyzed with two-way repeated measure ANOVA or three-way ANOVA as detailed in Supplemental Figure 2.

## RESULTS

### *Nalcn* transcripts are enriched in *Drd2*-SPNs

*Nalcn* gene product encodes a sodium leak channel that regulates the resting membrane potential and excitability of neurons. Although highly expressed in the striatum its function remains poorly characterized. By taking advantage of our striatal RNAseq database from *Drd2*-positives SPNs (Puighermanal et al., 2020), we found that *Nalcn* transcripts are enriched in the dorsal striatum (DS) and the nucleus accumbens (Acb) (comprising both shell and core) pellet fraction of *Drd2^Cre/+^;Ribotag^f/+^* mice compared with the input fraction (containing the mRNAs from all cellular types) (**Fig. 1a**). In addition, genes encoding its 3 ancillary subunits (*Unc-79*, *Unc-80*, *Fam155a*), required for its proper function and cellular localization were also found to be present in the DS and the Acb in both *Drd2*-positives and - negative SPNs (Puighermanal et al., 2020). Single molecule fluorescent *in situ* hybridization analysis confirmed the presence *Nalcn* transcripts in striatal *Drd2*-expressing SPNs of the DS and Acb in both males and females (**Fig. 1b**). Interestingly, *Nalcn* gene products were also detectable in *Drd1*-expressing SPNs, as well as cholinergic interneurons and striatal GABAergic interneurons whose identity remains to be determined (**Fig. 1b**). Together, these results indicate that *Nalcn* transcripts are expressed in striatal SPNs and interneurons.

**Figure 1.**
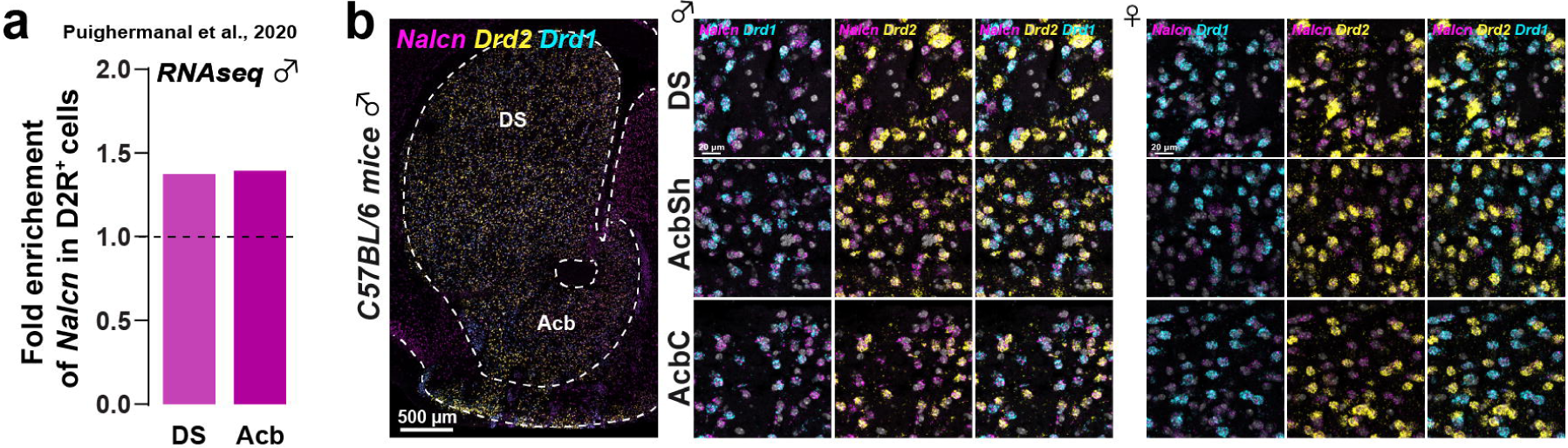
*Nalcn* expression in striatal SPNs. (**a**) Fold-change analysis showing enrichment of the *Nalcn* gene in the pellet fractions from the DS and Acb of *D2*-RiboTag mice (data extracted from Puighermanal et al. 2020). (**b**) Single-molecule fluorescent *in situ* hybridization of a coronal section from C57BL/6 mice stained with probes for *Nalcn* (magenta), *Drd2* (yellow) and *Drd1* (cyan) showing the DS and Acb regions (left panel). Higher magnification images show detailed views of the DS, AcbSh, and AcbC in male (middle panel) and female (right panel) of C57BL/6 mice.

### Generation of mice lacking NALCN in *Drd2*-SPNs

Because *Nalcn* transcripts are enriched in striatal *Drd2*-positives SPNs (**Fig. 1a**), we next investigated the functional role of the NALCN channel in *Drd2*-neurons. To do so, we generated conditional *Nalcn* knock-out mice expressing the calcium indicator GCamp6 in striatal *Drd2*-positives SPNs by crossing the *Drd2^Cre/+^* mouse line with the *Ai95^loxP^*;*Nalcn^loxP^*line (named thereafter *Drd2^Cre/+^*;*Ai95^f/+^*;*Nalcn^f/f^*) (**Supplemental Fig. 1**). As expected, GCamp6 expression was found in the DS and Acb (**Supplemental Fig. 1a upper left panel**) and detected in about half of striatal SPNs identified by STEP, a marker of both *Drd1*- and *Drd2*-expressing SPNs (**Supplemental Fig. 1a lower left panel**). Immunofluorescence and western blot analyses revealed no change in the expression levels of striatal proteins important for cell signaling including dopamine- and cAMP-regulated phosphoprotein, Mr = 32,000 (DARPP-32) (**Supplemental Fig 1b-c**), the glutamic acid decarboxylase 65- and 67- kilodalton isoforms (GAD65 and GAD67) (**Supplemental Fig 1c**), the vesicular inhibitory amino acid transporter (VGAT) (**Supplemental Fig. 1d**) as well as key component of the dopamine signaling namely the tyrosine hydroxylase (TH), the dopamine transporter (DAT) and the vesicular monoamine transporter 2 (VMAT2) (**Supplemental Fig. 2**). Finally, no change in the distribution of striatal GABAergic and cholinergic interneurons was detected in *Drd2^Cre/+^*;*Ai95^f/+^*;*Nalcn^f/f^*mice (**Supplemental Fig. 3**).

### Electrophysiological properties of striatal *Drd2*-SPNs lacking NALCN

Evidence indicate that NALCN channel is required for the pacemaker activity of spontaneously active neurons (Monteil et al., 2024; Ren, 2011), such as midbrain DA neurons (Philippart & Khaliq, 2018) or GABAergic neurons of the *substantia nigra pars reticulata* (Delgado-Zabalza et al., 2023; Lutas et al., 2016). Yet, SPNs are renowned for their highly hyperpolarized resting membrane potential and sparse spiking activity raising questions regarding the role of the NALCN in SPNs. To tackle this issue, we investigated the functional impact of NALCN in striatal *Drd2*-positive SPNs by using patch-clamp recording in striatal slices from male and female *Drd2^Cre/+^*;*Ai95^f/+^*(control) and *Drd2^Cre/+^*;*Ai95^f/+^*;*Nalcn^f/f^*(cKO) mice.

We first applied voltage steps from -100 to +80 mV to individual recorded neurons clamped at -70 mV. The calculated conductance at instantaneous current (**Fig. 2**) and steady state current (**Supplemental Fig. 4**) were significantly smaller for *Drd2*-positive SPNs recorded in cKO male mice compared to control male mice (**Fig. 2a** and **Supplemental Fig 4a**). In contrast, conductance and steady state current measured in *Drd2*-positive SPNs of female mice were not different between control and cKO mice (**Fig. 2b** and **Supplemental Fig 4b**) indicating that mechanisms may occur to compensate for the absence of NALCN channel in *Drd2*-positive SPNs of cKO female mice. Importantly, no differences were found between control and cKO in both males and females when similar analysis was performed in *Drd2*-negative SPNs (**Supplemental Fig 4c-d** and **Supplemental Fig. 5a-b**).

**Figure 2:**
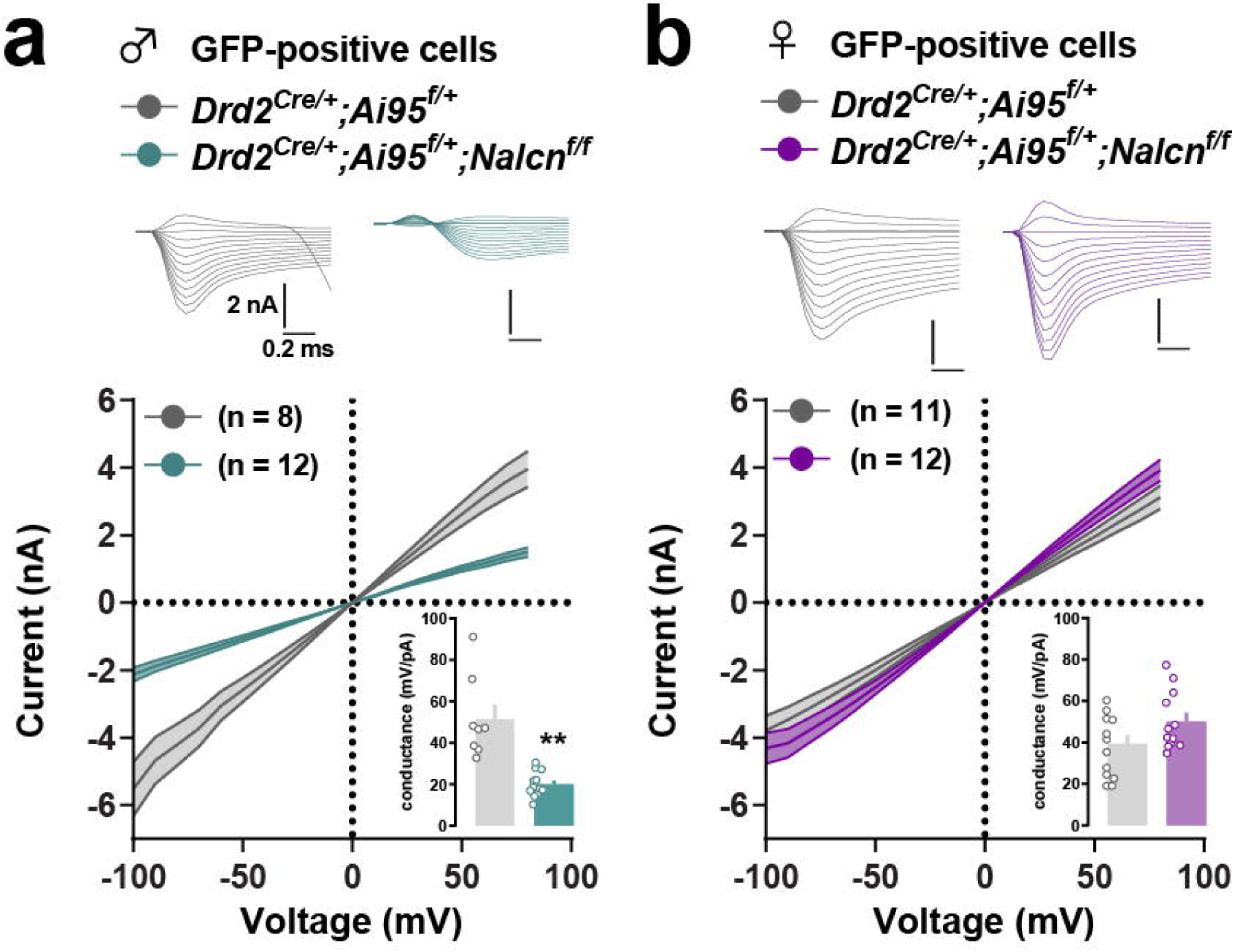
Instantaneous leak current in *Drd2*-positive SPNs of control and cKO mice. (**a-b**) Instantaneous current (nA) plotted as a function of applied voltage steps for *Drd2*-positive SPNs in male (**a**) and female (**b**) mice. Upper insets: illustrative traces of instantaneous leak current for voltage steps from -100 to +20 mV, 10 mV increment. Lower insets: conductance measured as the slope of the linear regression of individual curves. Values are represented as mean ± SEM. ** p < 0.01, (detailed statistical analysis in Supplemental Table 2: 2a-b).

The analysis of passive properties revealed that resting membrane potential of *Drd2*-positive SPNs was more depolarized in cKO male (**Fig. 3a**). The I/V curves showed an increased resistance in cKO male mice that was particularly noticeable for hyperpolarized steps (-150 to -130 pA) (**Fig. 3b-c**). Although female cKO mice displayed similar resting membrane potential changes than those observed in male cKO (**Fig. 3f**), resistance was not significantly different between control and cKO mice (**Fig. 3g-h**). Furthermore, rheobase was significantly increased in cKO female mice compared to control (**Fig. 3i-j**), which was not the case for male mice (**Fig. 3d-e**), supporting the hypothesis of compensatory mechanisms in female cKO mice. Finally, no difference in the resting membrane potential, resistance and rheobase were observed in *Drd2*-negative SPNs of males and females control compared to cKO mice (**Supplemental Fig. 6**)

**Figure 3:**
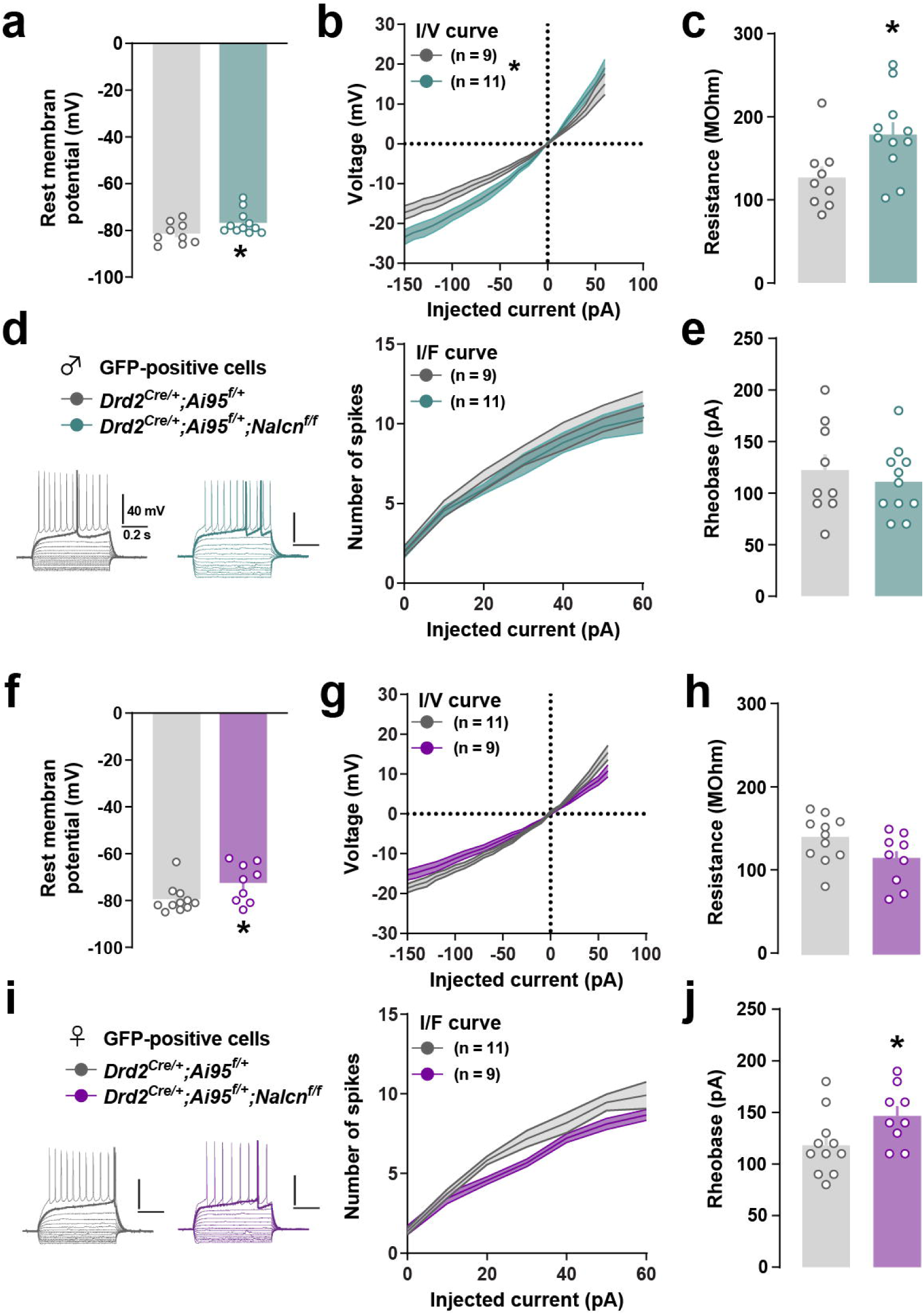
Intrinsic membrane electrophysiological properties of *Drd2*-positive SPNs in control and cKO mice. (**a** and **f**) Resting membrane potential (Rest memb potential) for *Drd2*-positive SPNs in male (**a**) and female (**f**) mice. (**b** and **g**) current/voltage curves for *Drd2*-positive SPNs in male (**b**) and female (**g**) mice. (**c** and **h**) resistance for *Drd2*-positive SPNs in male (**c**) and female (**h**) mice. (**d** and **i**) illustrative traces of voltage responses for current steps (600 ms) from -150 pA to +30 pA over rheobase, with an increment of 20 pA and current / number of spikes curves for *Drd2*-positive SPNs in male (**d**) and female (**i**) mice. (**e** and **j**) rheobase for *Drd2*-positive SPNs in male (**e**) and female (**j**) mice. Each dot represents one neuron. Values are represented as mean ± SEM. * p < 0.05 (detailed statistical analysis in Supplemental Table 2: 3a-j).

We next investigated the impact of *Nalcn* deletion in *Drd2*-positive SPNs on excitatory transmission in *Drd2*-positive and -negative SPNs in both male and female control and cKO mice (**Fig. 4** and **Supplemental Fig. 7**). Quantification of spontaneous EPSCs (sEPSCs) parameters revealed no difference for mean occurrence of events between control and cKO male mice (**Fig. 4a_1_**). Moreover, the amplitude of sEPSCs was significantly reduced and rise time significantly increased in *Drd2*-positive SPNs of cKO male mice (**Fig. 4a_2-3_**). In female mice, we noticed a trend to left-shift distribution of events amplitude (more events with small amplitude) and mean decay time of events was significantly increased (**Fig. 4b_1-3_**). No differences were found in *Drd2*-negative SPNs when males and females cKO mice were compared to their respective control, except for the distribution of inter-event intervals that was left-shifted in males, meaning more events with short intervals (**Supplemental Fig. 7a_1-3_-b_1-3_**). Of note, the paired-pulse ratio (50-ms interval) was not different between groups, suggesting an absence of alteration at the presynaptic level (**Supplemental Fig. 8a-d**).

**Figure 4:**
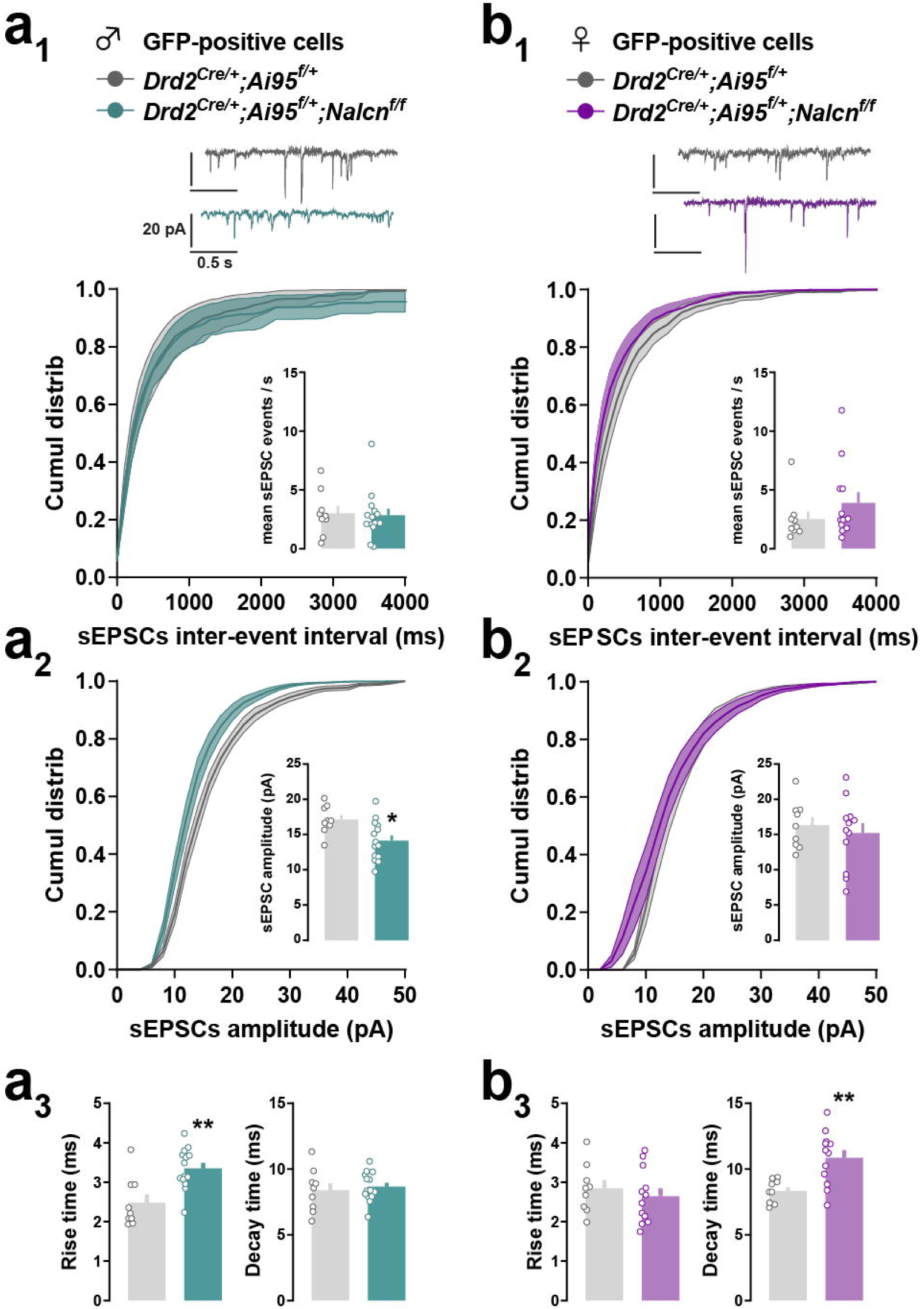
properties of sEPSCs events in *Drd2*-positive SPNs of control and cKO mice. (**a-b**) Properties of sEPSCs events recorded in *Drd2*-positive SPNs of male (**a**) and female (**b**) mice. (**a_1_-b_1_**) Cumulative distribution of sEPSCs inter-event intervals (ms) with mean number of events / s for each neuron in inset. (**a_2_-b_2_**) Cumulative distribution of sEPSCs amplitudes (pA) with mean amplitude for each neuron in inset. (**a_3_-b_3_**) Mean rise time (ms) (left) and decay time (ms) (right) of sEPSCs. Upper inset: illustrative trace of recordings for neurons clamped at -70 mV. Each dot represents the average value for one recorded neuron. Values are represented as mean ± SEM. * p < 0.05, ** p < 0.01, (detailed statistical analysis in Supplemental Table 2: 4a-b).

Finally, we tested whether the deletion of *Nalcn* in *Drd2*-positive SPNs alters the excitability of SPNs with EPSP/Spike coupling protocol that integrates both the membrane properties of the recorded neurons and its afferents (**Fig. 5**). Interestingly, while excitability was higher in *Drd2*-positive SPNs of male cKO mice compared to control mice, reduced excitability was observed in female cKO mice (**Fig. 5a-b**). The increased resistance and rheobase found previously in male and female cKO mice respectively, are likely to account for this opposing impact of *Nalcn* deletion on E/S coupling (**Fig. 3**). Surprisingly, similar changes were observed in *Drd2*-negative SPNs of male and female cKO mice suggesting that deletion of *Nalcn* in *Drd2*-positive SPNs alters local interactions between *Drd2*- and *Drd1*-positive SPNs (here identified as *Drd2*-negative SPNs) (**Supplemental Fig. 9a-b**). Together, these results suggest a sexual dimorphic effect of the invalidation of *Nalcn* in striatal *Drd2*-positive SPNs on passive membrane properties in *Drd2*-positive SPNs and on excitability in both *Drd2*- and *Drd1*-positive SPNs.

**Figure 5:**
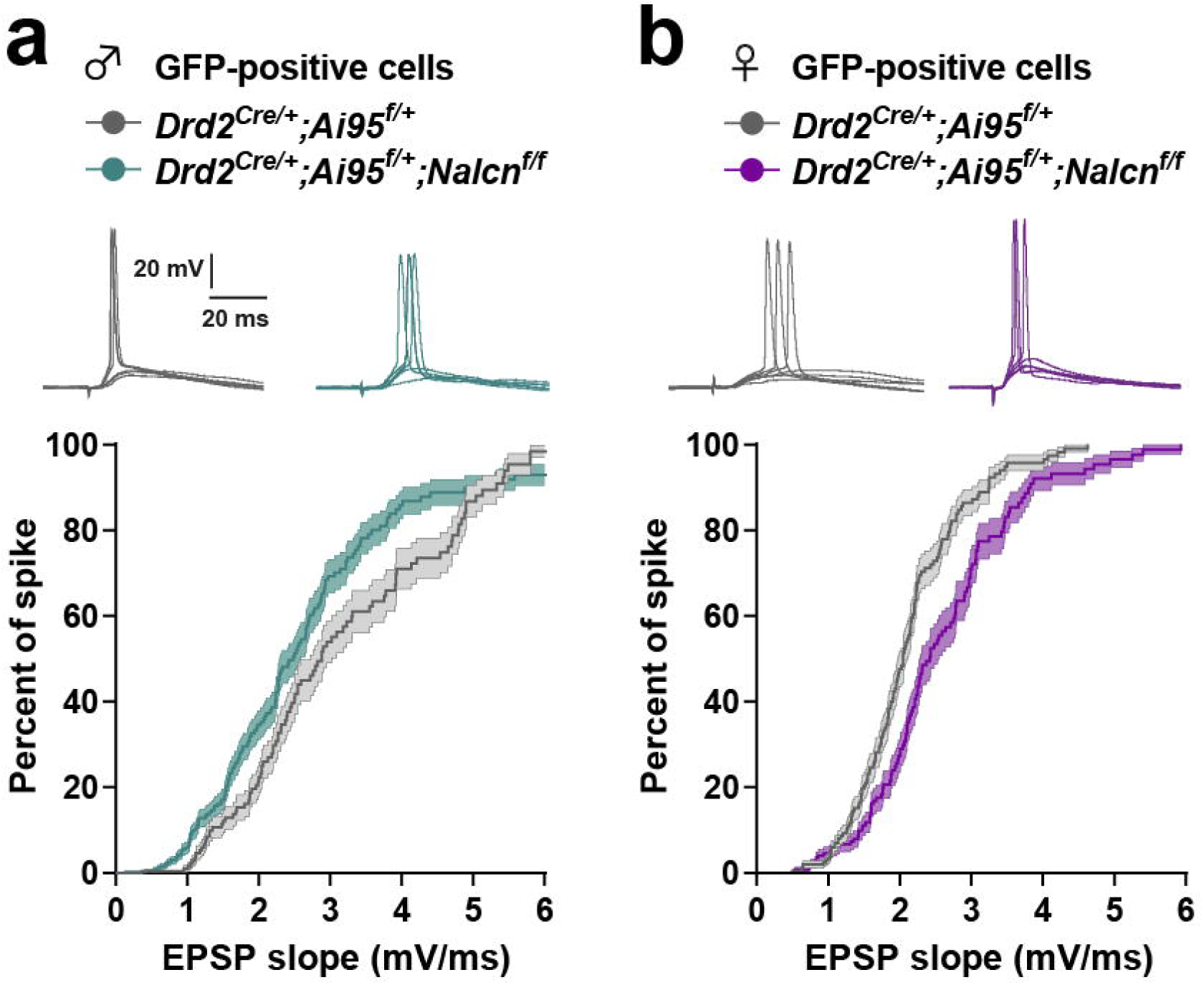
Excitability of *Drd2*-positive SPNs in control and cKO mice. (**a-b**) Survival curves of spike occurrence as a function of EPSP slope (mV/ms) for *Drd2*-positive SPNs in male (**a**) and female (**b**) mice. Upper insets: illustrative traces of neurons clamped at -60 mV with voltage responses for electrical stimulations of excitatory inputs with various intensities evoking EPSPs with or without spike. Values are represented as survival curves with mean ± SEM. (detailed statistical analysis in Supplemental Table 2: 5a-b).

### Eticlopride-induced intracellular signaling in *Drd2*-SPNs is enhanced female cKO mice

We then examined whether the deletion of *Nalcn* in *Drd2*-positive SPNs altered the ability of the selective D2R antagonist, eticlopride to regulate intracellular signaling in the dorsal striatum *Drd2*-positive SPNs. As expected, a significant increased number of pS6-235-236/cFOS and pS10-H3/cFOS immunoreactivity restricted to *Drd2*-SPNs was found in the dorsal striatum of both male and female control mice, 60 min after systemic administration of eticlopride (0.5 mg/kg) (**Fig. 6a-d**). In cKO male mice, eticlopride-induced pS6-S235/236, pS10-H3 and cFOS expression was similar to that observed in male control mice (**Fig. 6a-b**). In contrast, the number of pS6-235-236/cFOS and pS10-H3/cFOS-immunolabeled *Drd2*-SPNs was twofold higher in female cKO as compared to control mice (**Fig. 6c-d**). Of note, neither increased phosphorylation of pS6 and pH3 nor increased expression of cFOS were detected in *Drd2*-negative SPNs in either condition (**Fig. 6a-d**).

**Figure 6.**
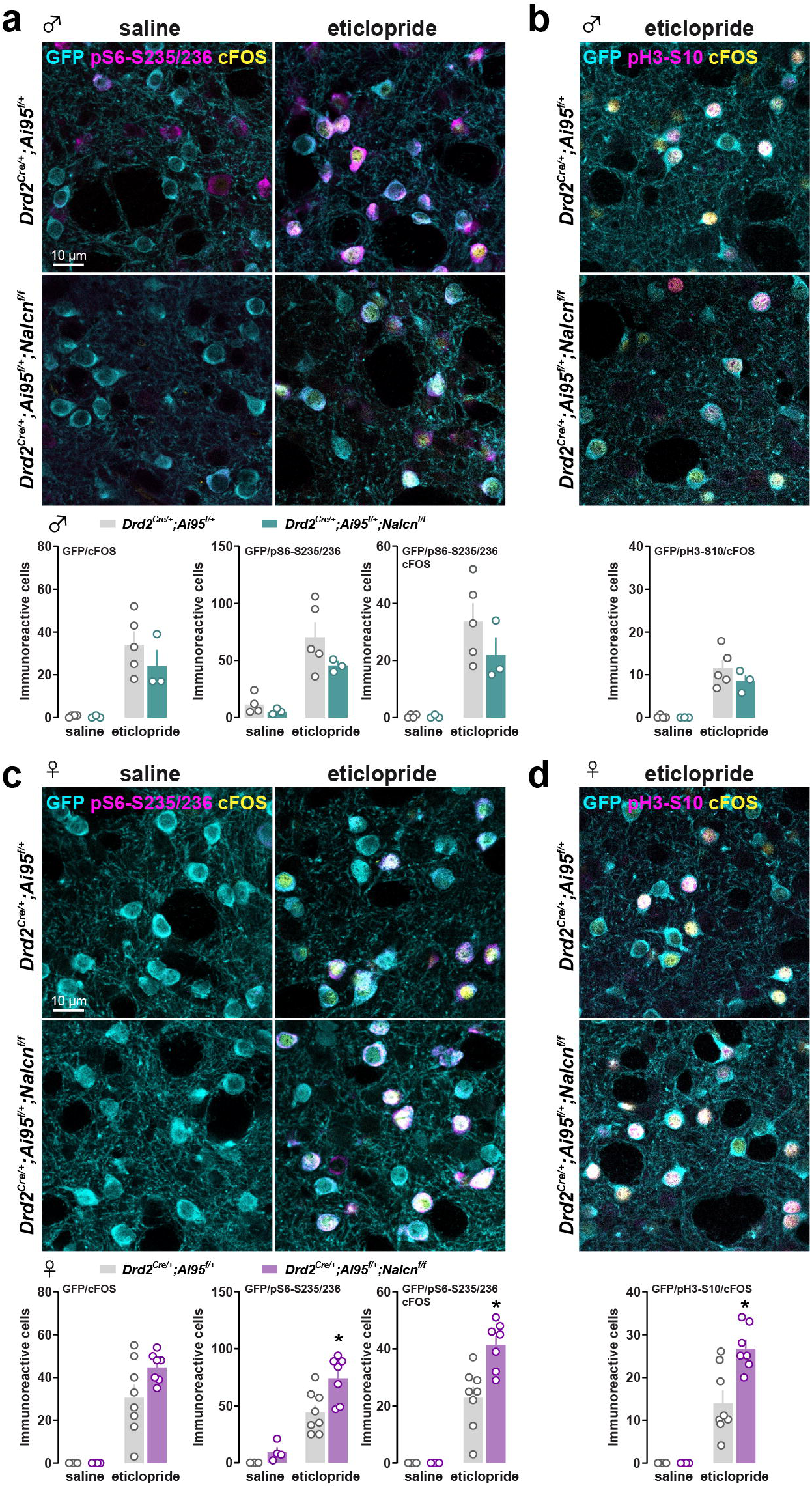
Eticlopride-induced changes in cellular reactivity in the DS. (**a** and **c**) Triple immunofluorescence staining for GFP (cyan), pS6-S235/236 (magenta) and cFOS (yellow) in the DS of male (**a**) and female (**c**) *Drd2^Cre/+^*;*Ai95^f/+^* (upper panels) and *Drd2^Cre/+^*;*Ai95^f/+^*;*Nalcn^f/f^*(lower panels) mice, 60 minutes after eticlopride or saline administration. Corresponding histograms illustrate the immunoreactive cells after saline or eticlopride administration. (**b** and **d**) Triple immunofluorescence staining for GFP (cyan), pH3-S10 (magenta) and cFOS (yellow) in the DS of male (**b**) and female (**d**) *Drd2^Cre/+^*;*Ai95^f/+^* (upper panels) and *Drd2^Cre/+^*;*Ai95^f/+^*;*Nalcn^f/f^* (lower panels) mice, 60 minutes after eticlopride administration. Histograms show the quantification of immunoreactive cells in each group. Values are represented as mean ± SEM. * p < 0.05 (detailed statistical analysis in Supplemental Table 2: a-d).

### Reduced food seeking behavior in male mice lacking *Nalcn* in *Drd2*-SPNs

Because *Drd2*-SPNs play an important role in motor control and motivation, we investigated whether any of these functions were altered in mice lacking *Nalcn* in *Drd2*-SPNs. We first assessed locomotion in a circular corridor. Male cKO mice displayed a slight basal horizontal and vertical locomotor hyperactivity as compared to control mice (**Fig. 7a, c**). This enhanced activity was not associated with altered motor coordination performance since no deficits were found in the beam walking test (**Supplemental Fig. 10a**). No alterations in locomotion and motor routines were observed between female mice lacking *Nalcn* in *Drd2*-SPNs and controls (**Fig. 7b, d** and **Supplemental Fig. 10b**). Finally, no differences were found in anxiety-like and repetitive behaviors assessed using the elevated plus maze (**Supplemental Fig. 11a-b**), marble burying test (**Supplemental Fig. 12a, c**) and digging behavior (**Supplemental Fig. 12b, d**).

**Figure 7.**
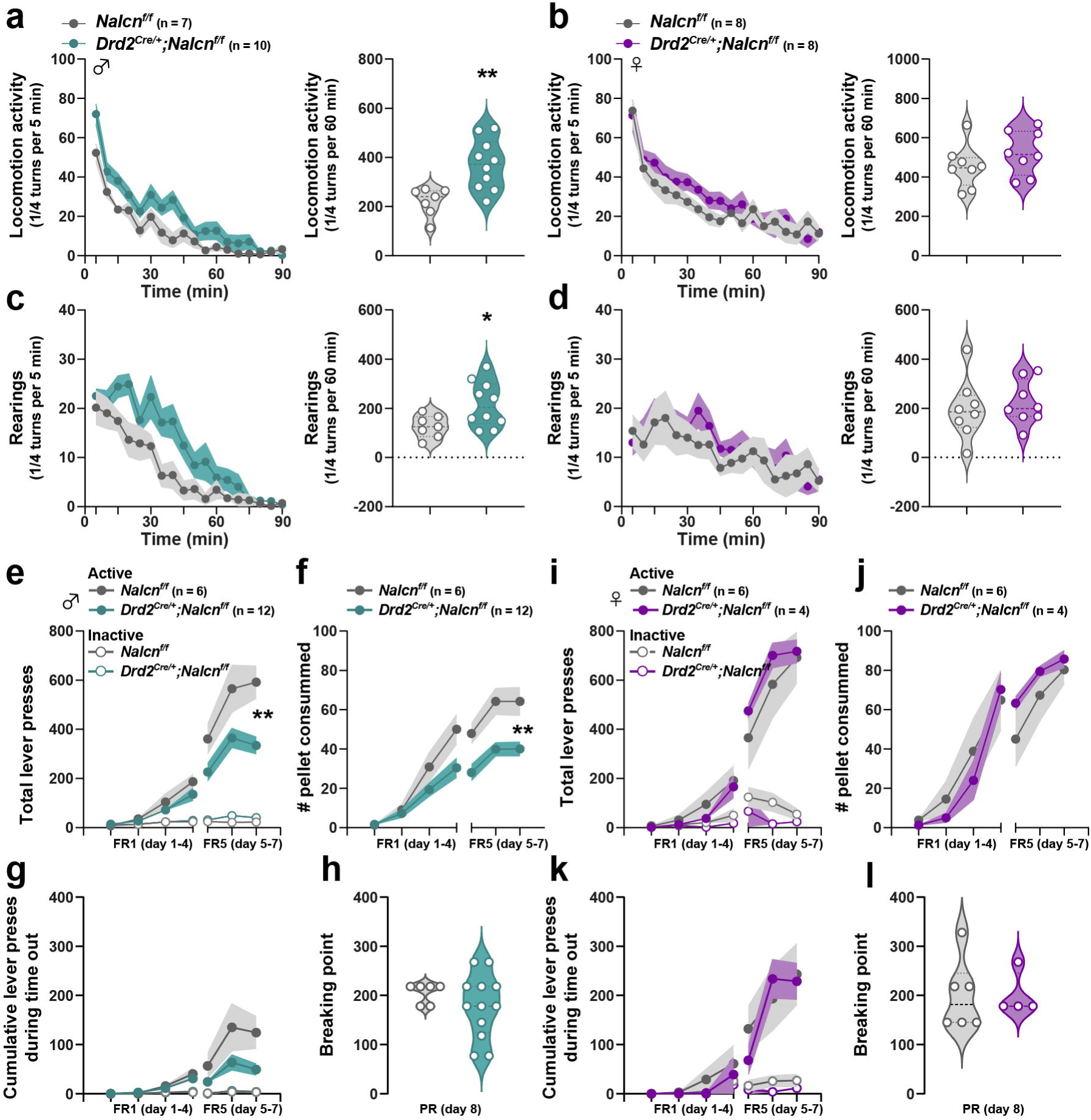
Behavioral assessment in control and cKO mice. (**a-b**) Horizontal locomotor activity measured in a circular corridor over 90 minutes in male (**a**) and female (**b**) *Nalcn^f/f^* and *Drd2^Cre/+^*;*Nalcn^f/f^*mice. (**c-d**) Vertical locomotor activity (rearing behavior) in male (**c**) and female (**d)** *Nalcn^f/f^* and *Drd2^Cre/+^*;*Nalcn^f/f^* mice. (**e** and **i**) Total lever presses for palatable food pellets on the active and inactive levers during FR1 (days 1-4) and FR5 (days 5-7) in male (**e**) and female (**i**) *Nalcn^f/f^* and *Drd2^Cre/+^*;*Nalcn^f/f^* mice. (**f** and **j**) Daily pellet consumption during FR1 and FR5 in male (**f**) and female (**j**) *Nalcn^f/f^* and *Drd2^Cre/+^*;*Nalcn^f/f^* mice. (**g** and **k**) Cumulative lever presses during time-out periods on the active and inactive levers in male (**g**) and female (**k**) *Nalcn^f/f^* and *Drd2^Cre/+^*;*Nalcn^f/f^*mice during FR1 and FR5. (**h** and **l**) Breaking point analysis under a PR reinforcement schedule in male (**h**) and female (**l**) *Nalcn^f/f^*and *Drd2^Cre/+^*;*Nalcn^f/f^* mice. * p < 0.05, ** p < 0.01 (detailed statistical analysis in Supplemental Table 2: 7a-l).

We then investigated motivation by assessing reward-seeking behaviors. Food-restricted male and female control and cKO mice were trained on a fixed-ratio 1 (FR1) reinforcement schedule for isocaloric palatable food delivery for 4 consecutive days. Under FR1, similar increased total number of active lever presses and pellets consumed were observed between control and cKO male mice (**Fig. 7e-f**). As the cost of reward increased (FR5 schedule), cKO male mice significantly reduced their number of active lever presses and consumed pellets compared to control mice suggesting that goal-oriented motivation might be altered (**Fig. 7e-f**). This difference was not the result of greater response disinhibition since no difference in premature responses on the active lever during the time-out period was observed between the control and cKO mice (**Fig. 7g**). To probe whether willingness and/or vigor for food-seeking behavior was altered in cKO male mice, we analyzed lever presses under a progressive ratio (PR) schedule of reinforcement (**Fig. 7h**). Although breaking points were not significantly different, the variability was much higher in cKO than in control mice as illustrated by the significant difference in variances, suggesting that goal-oriented motivation was altered in cKO male mice (**Fig. 7h**). In contrast, while 3 control and 4 cKO female mice never learnt the task and were excluded from the analysis, the remaining control and cKO female mice similarly increased their total number of active lever presses and consumed pellets under under FR1 and FR5 (**Fig. 7i-k**) and displayed similar breaking points (**Fig. 7l**). Altogether, these results indicate that altered NALCN signaling in *Drd2*-SPNs reduced food-seeking behaviors in male mice.

### Altered discriminative avoidance behavior in male mice lacking *Nalcn* in *Drd2*-SPNs

Finally, we evaluated whether altered NALCN functions in *Drd2*-SPNs impact learned defensive behaviors. We first tested the ability to discriminate between auditory signals indicating (CS+) or not (CS-) that engaging shuttling from one compartment to another prevent the onset of a threat (**Fig. 8a**). With time, both control and cKO male mice similarly increased their avoidance probability following the presentation of the CS+ (**Fig. 8b**). However, while control male mice displayed a low probability of defensive responses following CS-presentation, these responses gradually increased in cKO male mice leading to weak discriminative performance compared to control mice (**Fig. 8c-d**).

**Figure 8.**
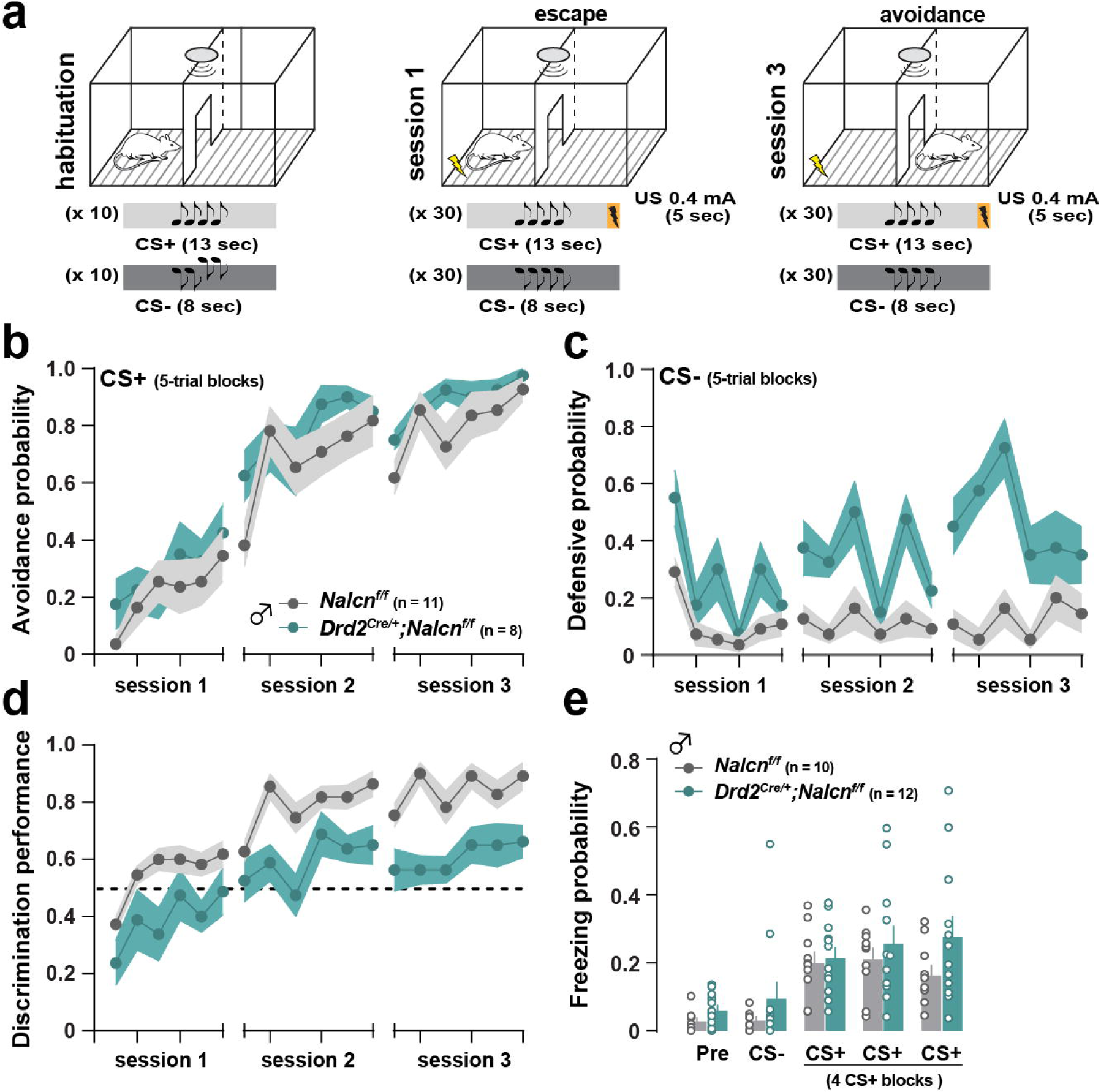
**Discriminative aversive behavior in control and cKO mice**. (**a**) Schematic cartoon describing the protocol used to evaluate discriminative active avoidance learning. (**b**) Avoidance probability measured as the frequency of movements to the neighboring compartment during the CS+ presentation among 3 sessions in male *Nalcn^f/f^*and *Drd2^Cre/+^*;*Nalcn^f/f^* mice. (**c**) Defensive probability indicating the rate at which the animal exhibited an escape or avoidance response during the CS-presentation across 3 sessions in male *Nalcn^f/f^* and *Drd2^Cre/+^*;*Nalcn^f/f^*mice. (**d**) Discrimination performance between CS+ and CS-in male *Nalcn^f/f^* and *Drd2^Cre/+^*;*Nalcn^f/f^*mice. (**e**) Freezing probability in mice that underwent discriminatory fear conditioning before (pre) and after the presentation of CS+ and CS-in male *Nalcn^f/f^* and *Drd2^Cre/+^*;*Nalcn^f/f^* mice. ** p < 0.01 (detailed statistical analysis in Supplemental Table 2: 8b-e).

To determine whether impaired discriminative performance observed in cKO male mice results from general altered discriminative learnings associated with threat, we analyzed freezing responses in mice that underwent discriminative auditory fear conditioning. Following the day of conditioning, both control and cKO male mice displayed higher probability of freezing responses following CS+ presentation compared to CS-presentation (**Fig. 8e**). Together, our results suggest that exacerbated avoidance probability observed in cKO mice during CS-presentation did not result from general impairment of discriminative learning associated with a threat.

## DISCUSSION

The present study unveils a previously undescribed sexual dimorphic effect of *Nalcn* invalidation in *Drd2*-expressing cells on striatal-dependent physiology, signaling and behaviors. Indeed, while passive properties of *Drd2*-positive SPNs in male cKO mice are profoundly altered, only few parameters are modified in female. Yet, excitability of both *Drd2*- and *Drd1*-SPNs is modified by selective NALCN invalidation in *Drd2*-SPNs, in both male and female cKO mice. In contrast, eticlopride-induced intracellular signaling in *Drd2*-SPNs is enhanced only in female cKO mice. Finally, we found that NALCN in *Drd2*-SPNs regulates locomotion, reward-seeking and dynamic defensive behaviors only in male mice.

Physiology of NALCN channels has been extensively studied in cultured neurons, with precise pharmacology and modulation of ions concentrations in extra- and intra-cellular compartments (Lu et al., 2007; Ren, 2011). In more integrative models, evidence indicate that NALCN conductance participates to resting membrane potential, pacemaker activity and excitability of dopaminergic neurons from the *substantia nigra pars compacta* (Philippart & Khaliq, 2018; Um et al., 2021) and the ventral tegmental area (Cobb-Lewis et al., 2023), as well as GABAergic neurons from the *substantia nigra pars reticulata* (Delgado-Zabalza et al., 2023; Lutas et al., 2016). Beyond its sodium leak activity, NALCN function is tightly controlled by its associated proteins, forming a channelosome (Monteil et al., 2024), by extracellular calcium and magnesium concentrations (Chua et al., 2020; Ren, 2011) and is strongly modulated by G protein-coupled receptors activated by hormones and neuromodulators (Monteil et al., 2024). Here, we assessed the impact of NALCN deletion in *Drd2*-expressing neurons on membrane properties and excitability of SPNs in acute brain slices of both adult male and female mice. These analyses conducted without any pharmacology or modification of ion concentrations revealed several unexpected results. At first, resting membrane potential is more depolarized in *Drd2*-SPNs of male and female cKO mice, which is opposite to what is observed generally when NALCN channel is blocked in neurons (Monteil et al., 2024; Philippart & Khaliq, 2018; Ren, 2011). Considering that NALCN channels are involved in the balance between potassium and sodium basal permeability of neurons to adjust their resting membrane potential (Ren, 2011), one can hypothesize that this counterintuitive effect in SPNs is due to a compensation of the system with a down-regulation of potassium channels in *Drd2*-SPNs of cKO mice. Alternatively, a functional coupling between NALCN and Na^+^-activated K^+^ channels could account for the more depolarized RMP observed in the present study. Indeed, recent evidence from myometrial cells indicate that the negative RMP of these cells relied in part of the functional coupling between NALCN and the Slo2.1 Na^+^-activated K^+^ channel (Ferreira et al., 2021, 2024). Given the enrichment in *Drd2*-SPNs of *Kcnt1* transcripts encoding the Slo2.2 channel, another related Na^+^-activated K^+^ channel (Puighermanal et al., 2020), future experiments will determine whether functional coupling between NALCN and this Na^+^-activated K^+^ channels is at play to regulate RMP in SPNs.

Furthermore, our results suggest that the strategies for compensation in male and female mice are different. Indeed, in males, *Drd2*-SPNs membrane resistance is significantly increased, which is in accordance with the absence of NALCN channels and the putative reduction of potassium channels. In contrast, in female, the resistance of *Drd2*-SPNs is unaffected and tend to be decreased, while rheobase is significantly increased. As a consequence, excitability as measured by the E/S coupling protocol, is increased in males while it is decreased in females. Although not tested in the present study, compensatory expression of the nonselective cation channels TRPC3 in females could account for such differences, as recently demonstrated in DA neurons of the *substantia nigra pars compacta*, though sex was not considered as biological variable (Um et al., 2021). Finally, our recordings conducted in both male and female mice revealed that NALCN deletion in *Drd2*-SPNs do not affect membrane properties of *Drd1*-SPNs attesting the specificity of NALCN invalidation and the lack of cell non-autonomous effect. Yet, excitability of *Drd1*-SPNs in cKO mice is increased in males and decreased in females, similar to what is observed in *Drd2*-SPNs. Although additional work is required to uncover the underlying mechanisms, these results further illustrate the importance of local interactions between *Drd2*- and *Drd1*-SPNs in the homeostasis of striatal physiology (Burke et al., 2017; Taverna et al., 2008).

Mounting evidence suggest that NALCN channels may act as an effector of G protein-coupled receptors to modulate, either positively or negatively, neuronal activity (Monteil et al., 2024). Recently, the existence of a functional coupling between NALCN and D2R has been unveiled. Thus, inhibition of NALCN following D2R activation has been proposed to be a core mechanism through which D2R control midbrain DA neurons excitability (Cobb-Lewis et al., 2023; Philippart & Khaliq, 2018). Given the high expression of NALCN in *Drd2*-SPNs, we thought to probe the impact of the deletion of NALCN on D2R-mediated intracellular signaling. In the striatum, activation of intracellular signaling in *Drd2*-SPNs can be assessed by administering D2R antagonist such as raclopride or eticlopride (Valjent et al., 2019). We found that phosphorylation of histone H3 and ribosomal protein S6 as well as cFOS expression induced by eticlopride were twofold increased in female cKO compared to control mice suggesting that in female but not in male, NALCN expressed by *Drd2*-SPNs might act as a brake preventing exacerbated intracellular events in response to D2R blockade. The absence of altered striatal levels of *Drd2* transcripts and D2R protein expression between male and female of both genotypes suggests that the observed effects most likely result from a dysfunctional coupling between NALCN and D2R. Although not yet established, such disruption of NALCN/D2R coupling could impact adenosine A2 receptor (A2aR) signaling which has been shown to be critical to mediate the activation of intracellular signaling in *Drd2*-SPNs in response to D2R blockage (Bertran-Gonzalez et al., 2009; Valjent et al., 2011). Whether NALCN act as a direct or indirect effector of A2aR to modulate the activity of *Drd2*-SPNs remains to be established.

Our study unveils that in male but not female, the loss of NALCN in *Drd2*-SPNs resulted in an increased locomotor activity. This phenotype, reminiscent to the one observed in mice bearing a deletion of *Ppp1r1b* encoding DARPP-32 (Bateup et al., 2010) or the fat mass and obesity associated gene (*Fto*) (Ruud et al., 2019) selectively in *Drd2*-SPNs, suggesting that similarly to DARPP-32 and FTO, NALCN function contributes to the basal inhibitory tone on locomotion exerted by *Drd2*-SPNs. It is important to note that we found no alteration of motor routines, perseveration and coordination despite the fact that *Nalcn* is most likely invalidated in Purkinje cells as a consequence of the Cre recombinase expression under the transcriptional control of the *Drd2* promotor (Cutando et al., 2022). Although altered functions of Drd2-SPNs have been associated to impaired operant conditioning (Augustin et al., 2020), the impact of loss of NALCN in Drd2-SPNs on goal-directed behaviors appears marginal. Indeed, mice lacking *Nalcn* in *Drd2*-expressing cells rapidly acquired instrumental learning under FR1 but as the operant efforts to obtain the reward increased, cKO male but not female mice reduced their active lever presses suggesting a decrease of motivation or/and vigor for food-seeking behavior. However, despite no difference was found under progressive ratio, breaking point values were more variable in cKO male mice suggesting that functional NALCN in Drd2-SPNs might play an important role in the regulation of the motivational drive depending on the metabolic state. Future experiments are needed to determine whether as recently exemplified for *Drd2* signaling in WFS1-neurons (Castell et al., 2024), NALCN in *Drd2*-SPNs regulate homeostatic-dependent food operant behavior.

Given the prominent role of *Drd2*-SPNs in the regulation of defensive responses (Nguyen et al., 2019), we also assessed whether reactive and passive defensive behaviors towards auditory cues predicting threat were impacted by the loss of NALCN in *Drd2*-expressing cells. Using a discriminative active avoidance paradigm, we found that male mice lacking *Nalcn* in *Drd2*-expressing cells efficiently switched from escape to avoidance responses in presence of an auditory threat-predicting cue. This indicates that, even if *Nalcn* is invalidated in *Drd2*-expressing of the central nucleus of the amygdala and the bed nucleus of the stria terminalis (De Bundel et al., 2016), associative learning associated with negative valence was not impaired. However, male cKO mice also exhibited strong defensive responses towards safety auditory cues. Although such low discriminative performance may suggest excessive generalization of threat responses (De Bundel et al., 2016; Dunsmoor & Paz, 2015), several lines of evidence argue against this hypothesis. Thus, cKO mice do not exhibit high level of anxiety normally associated with overgeneralization (Lissek et al., 2005) as approach-avoidance behaviors were comparable to control mice. Moreover, acquisition, consolidation and expression of discriminative learning between stimuli representing threat or safety appear not to be impacted by the loss of NALCN in *Drd2*-expressing cells since cKO mice displayed higher levels of freezing responses following CS+ presentation compared to CS-. Although we cannot exclude that the increased probability of defensive responses during CS-in active avoidance may directly result from the basal hyperlocomotor phenotype, future studies will determine whether other functions such as the recruitment of attentional resources or the selection of appropriate defensive behavioral responses are impacted by the loss of NALCN in *Drd2*-expressing cells.

It is important to keep in mind that using the Drd2-cre driver line and given the widespread expression of NALCN in brain, its invalidation is not restricted to Drd2-SPNs therefore precluding to establish strict causality between altered electrophysiological properties of *Drd2*-positive SPNs and behavioral phenotypes. Indeed, although motor control, food-seeking and defensive behaviors may involve striatal *Drd2*-expressing cells (De Bundel et al., 2016; Mourra et al., 2020; Olivetti et al., 2024; Trifilieff et al., 2013; Walle et al., 2024), we cannot exclude that invalidation in other cell-type account for some of the phenotypical traits. For instance, NALCN channels are highly expressed in midbrain DA neurons (Cobb-Lewis et al., 2023; Lutas et al., 2016) and striatal cholinergic interneurons, two neuronal population targeted using the *Drd2*-cre driver line. Future experiments are required to determine whether selective deletion of *Nalcn* within DA neurons and/or cholinergic interneurons phenocopies basal hyperactivity and altered food-seeking behaviors observed in our cKO mice.

## Acknowledgements

The authors thank Pr Dejian Ren for generously providing the *Nalcn^f/f^* mice. We thank MRI platform (Biocampus). The authors also thank iExplore from IGF for their involvement in the maintenance and breeding of the colonies. This work was supported by CNRS, INSERM, Fondation pour la Recherche Médicale (EQU202203014705, EV), the French National Research Agency (ANR-20-CE14-0020 and ANR-21-CE16-0028 to EV, ANR-21-NEU2-0004-01 in the frame of the the ERA-NET Neuron 2021 call on neurodevelopmental disorders (https://www.neuron-eranet.eu/projects/RestoreLeak/), AM), the Labex “Ion Channel Science and Therapeutics” (ANR-11-LABX-0015, AM).

## Author contribution

L.C., F.B., C. B.B. and E.V. conceived the study and designed the experiments. L.C. performed biochemical and behavioural experiments. A.G performed behavioural experiments. C. B.B. designed, performed and analyzed *ex vivo* patch-clamp recordings. C.N., M.A. and A.M. performed immunostaining experiments. L. M. performed the fluorescent *in situ* hybridization. C. B. performed genotyping. P. L. and A.M. generated double and triple transgenic mice. L.C., A.M., C. B.B. and E.V. wrote the manuscript designed the study with inputs from all authors.

