## Supplemental material for "Sex-biased effect of sodium leak channel NALCN deletion in striatal *Drd2* spiny projection neurons"

### Supplemental Figures

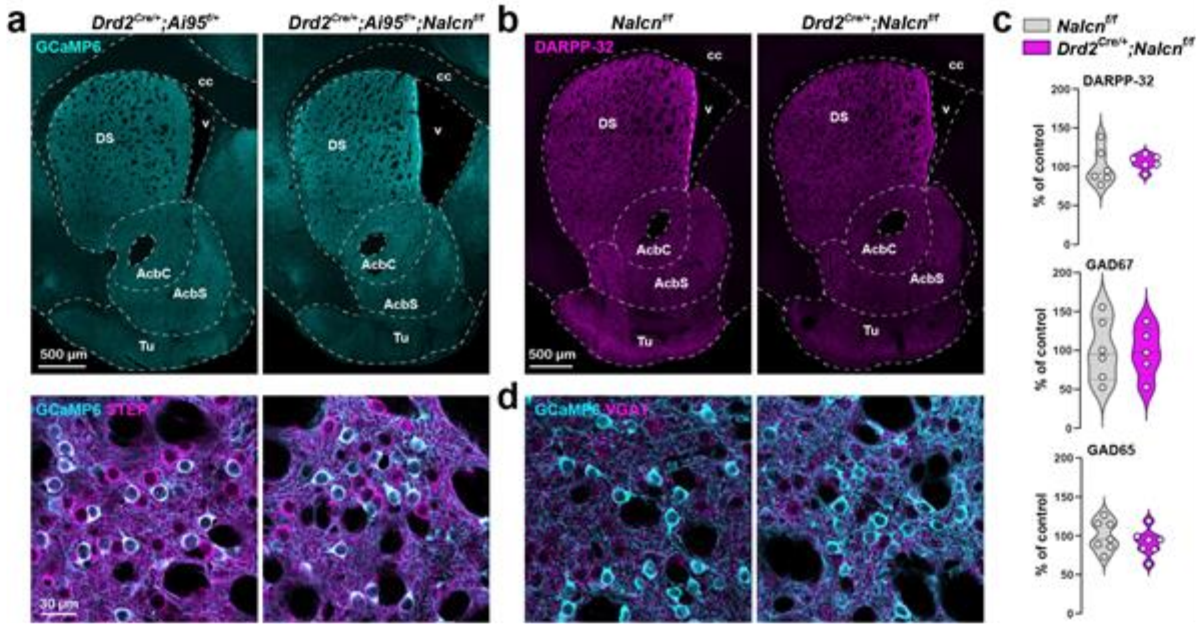

**Supplemental Figure 1: Biochemical characterization of *$Drd2^{Cre/+}$;$Ai95^{f/+}$;$Nalcn^{f/f}$* and *$Drd2^{Cre/+}$;$Nalcn^{f/f}$* mice.** (**a**) Immunofluorescence staining for GCaMP6 (cyan) (upper panel) and GCaMP6 (cyan) and STEP (magenta) (lower panel) in coronal sections of *$Drd2^{Cre/+}$;$Ai95^{f/+}$* (left) and *$Drd2^{Cre/+}$;$Ai95^{f/+}$;$Nalcn^{f/f}$* (right) mice. (**b**) Immunofluorescence staining for DARPP-32 in coronal sections of *$Nalcn^{f/f}$* (left) and *$Drd2^{Cre/+}$;$Nalcn^{f/f}$* (right) mice. (**c**) Western blot quantification of striatal DARPP-32, GAD67 and GAD65 expression in *$Nalcn^{f/f}$* and *$Drd2^{Cre/+}$;$Nalcn^{f/f}$* mice. (**d**) Double immunofluorescence staining for striatal GCaMP6 (cyan) and VGAT (magenta) in *$Nalcn^{f/f}$* (left) and *$Drd2^{Cre/+}$;$Nalcn^{f/f}$* (right) mice (detailed statistical analysis in Supplemental Table 2: S1c).

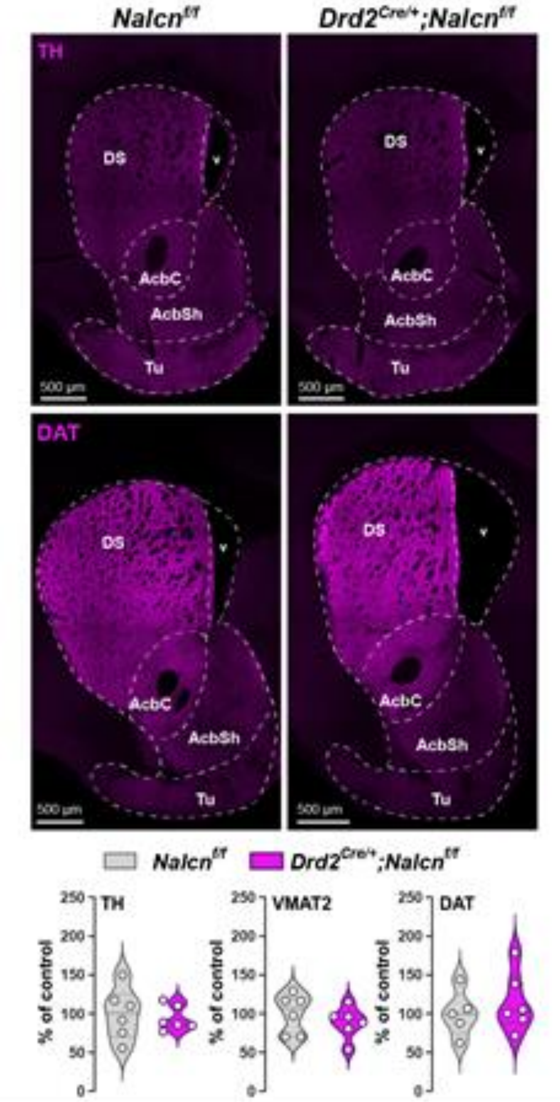

**Supplemental Figure 2: Biochemical characterization of *Drd2$^{Cre/+}$;Nalcn$^{f/f}$* mice.** (**a**) Immunofluorescence staining for TH (upper panels) and DAT (lower panels) in coronal sections of *Nalcn$^{f/f}$* (left) and *Drd2$^{Cre/+}$;Nalcn$^{f/f}$* (right) mice. (**b**) Western blot quantification of striatal TH, VMAT2 and DAT expression in *Nalcn$^{f/f}$* (control) and *Drd2$^{Cre/+}$;Nalcn$^{f/f}$* (cKO) mice (detailed statistical analysis in Supplemental Table 2: S2).

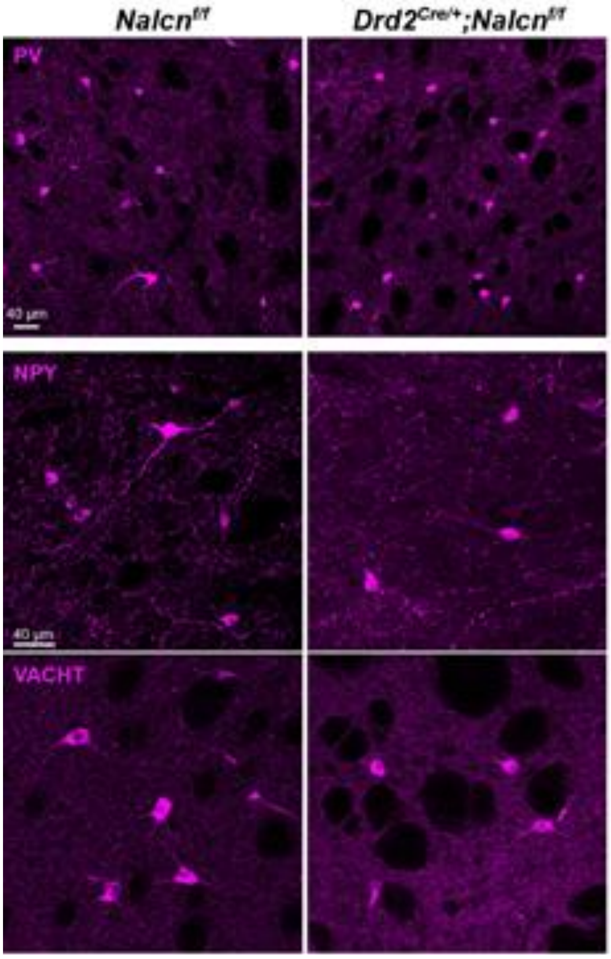

**Supplemental Figure 3: Distribution of GABAergic and cholinergic interneurons in *Drd2$^{Cre/+}$;Nalcn$^{f/f}$* mice.** Immunohistochemistry staining of striatal PV (upper panel), NPY (middle panel) and VACHT (lower panel) in *Nalcn$^{f/f}$* (control) and *Drd2$^{Cre/+}$;Nalcn$^{f/f}$* (cKO) mice.

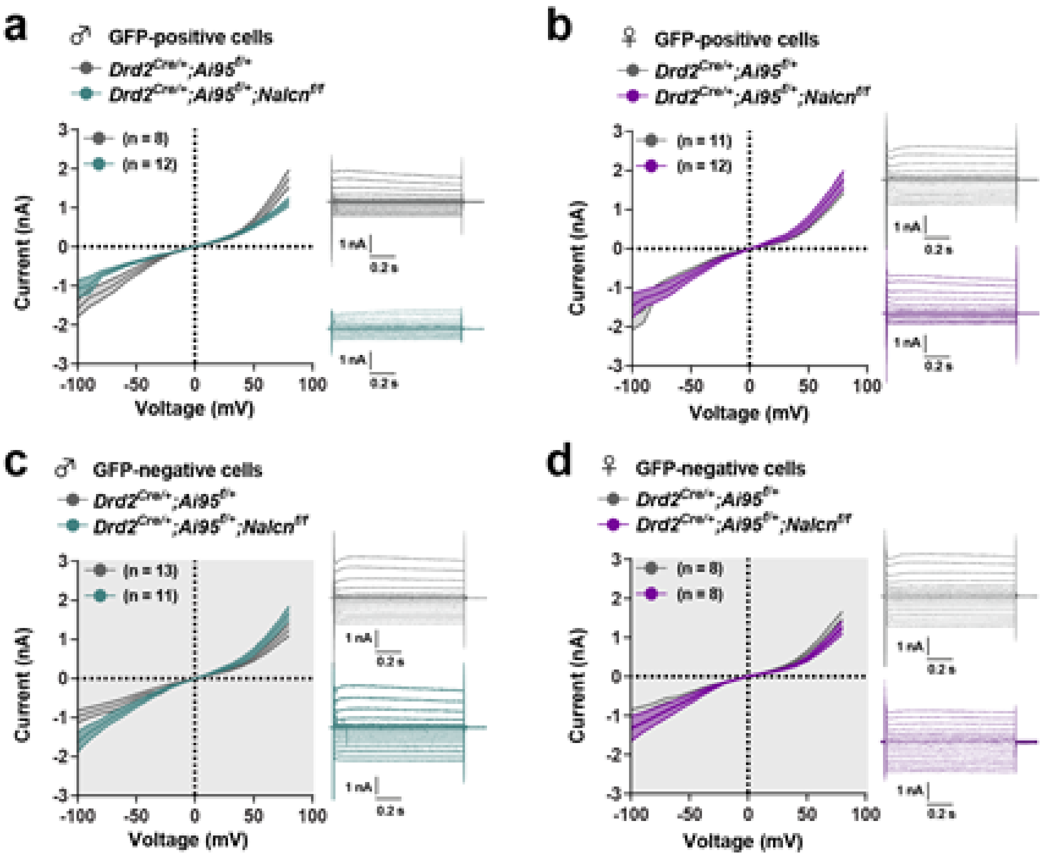

**Supplemental Figure 4: Steady-state leak current in SPNs of control and cKO mice.** (**a-d**) Steady-state current (nA) plotted as a function of applied voltage steps for *Drd2*-positive SPNs in male (**a**) and female (**b**) mice and for *Drd2*-negative SPNs in male (**c**) and female (**d**) mice. Right insets: illustrative traces of steady-state leak currents for voltage steps from -100 to +80 mV with 10 mV increment. Values are represented as mean ± SEM. (detailed statistical analysis in Supplemental Table 2: S4a-d).

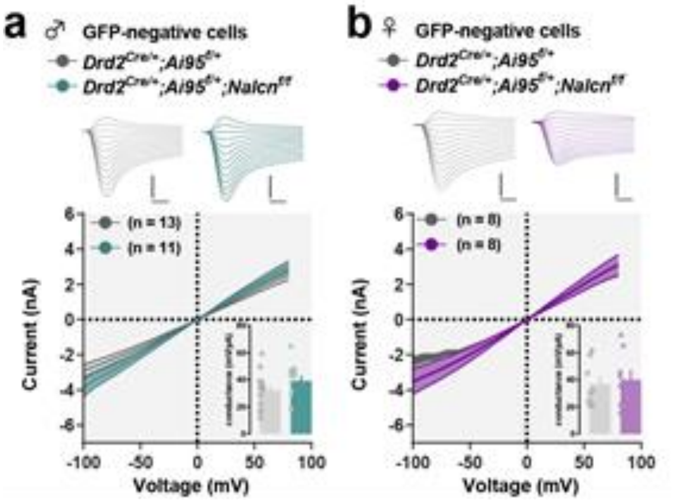

**Supplemental Figure 5: Instantaneous leak current in *Drd2*-negative SPNs of control and cKO mice.** (**a-b**) Instantaneous current (nA) plotted as a function of applied voltage steps for *Drd2*-negative SPNs in male (**a**) and female (**b**) mice. Upper insets: illustrative traces of instantaneous leak current for voltage steps from -100 to +20 mV, 10 mV increment. Lower insets: conductance measured as the slope of the linear regression of individual curves. Values are represented as mean ± SEM. (detailed statistical analysis in Supplemental Table 2: S5a-b).

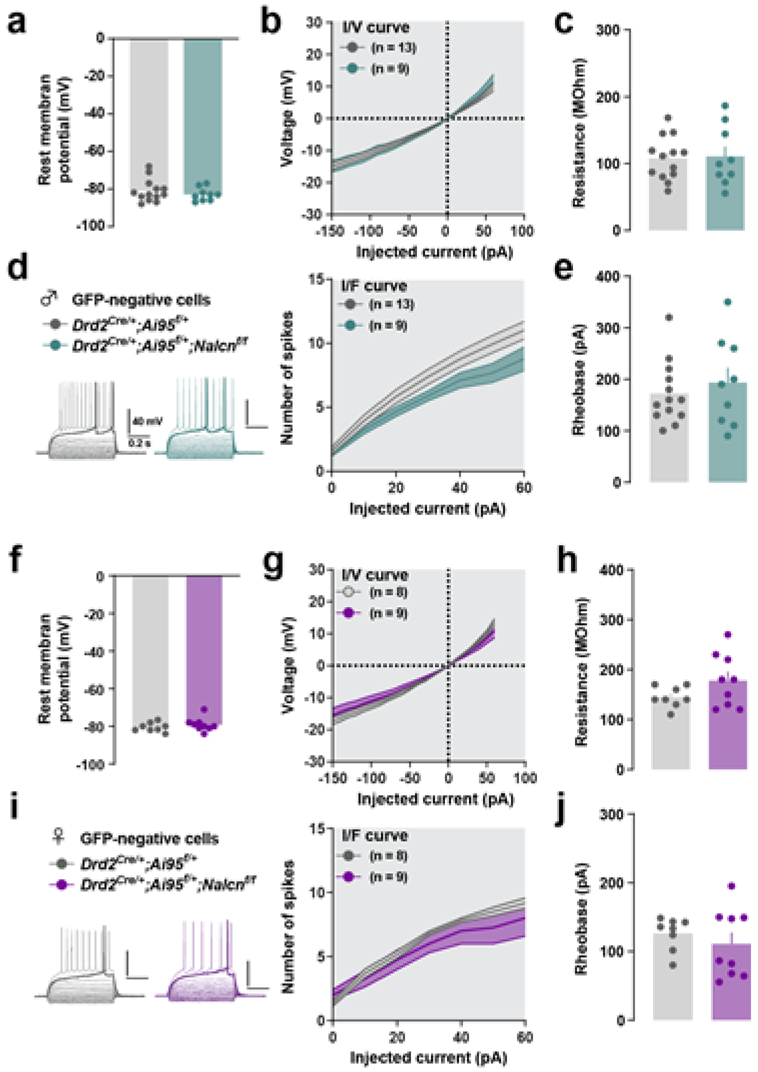

**Supplemental Figure 6: Intrinsic membrane electrophysiological properties of *Drd2*-negative SPNs in control and cKO mice.** (**a** and **f**) Resting membrane potential for *Drd2*-positive SPNs in male (**a**) and female (**f**) mice. (**b** and **g**) current/voltage curves for *Drd2*-positive SPNs in male (**b**) and female (**g**) mice. (**c** and **h**) resistance for *Drd2*-positive SPNs in male (**c**) and female (**h**) mice. (**d** and **i**) illustrative traces of voltage responses for current steps (600 ms) from -150 pA to +30 pA over rheobase, with an increment of 20 pA and current / number of spikes curves for *Drd2*-positive SPNs in male (**d**) and female (**i**) mice. (**e** and **j**) rheobase for *Drd2*-positive SPNs in male (**e**) and female (**j**) mice. Each dot represents one neuron. (detailed statistical analysis in Supplemental Table 2: S6a-j).

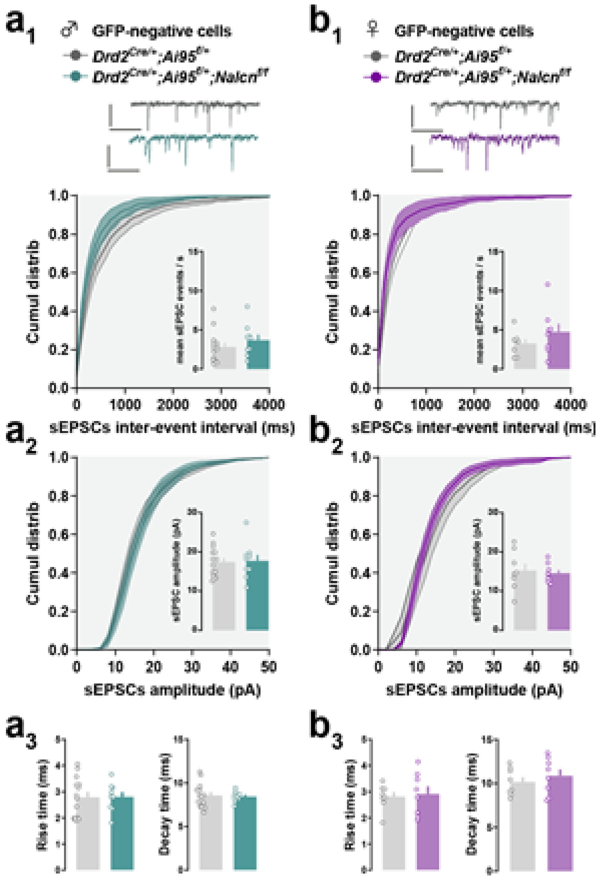

**Supplemental Figure 7: properties of sEPSCs events in *Drd2*-negative SPNs of control and cKO mice.** (**a-d**) (**a-b**) Properties of sEPSCs events recorded in *Drd2*-negative SPNs of male (**a**) and female (**b**) mice. (**$a_1$-$b_1$**) Cumulative distribution of sEPSCs inter-event intervals (ms) with mean number of events / s for each neuron in inset. (**$a_2$-$b_2$**) Cumulative distribution of sEPSCs amplitudes (pA) with mean amplitude for each neuron in inset. (**$a_3$-$d_3$**) Mean rise time (ms) (left) and decay time (ms) (right) of sEPSCs. Upper inset: illustrative trace of recordings for neurons clamped at -70 mV. Each dot represents the average value for one recorded neuron. Values are represented as mean ± SEM. (detailed statistical analysis in Supplemental Table 2: S7a-b).

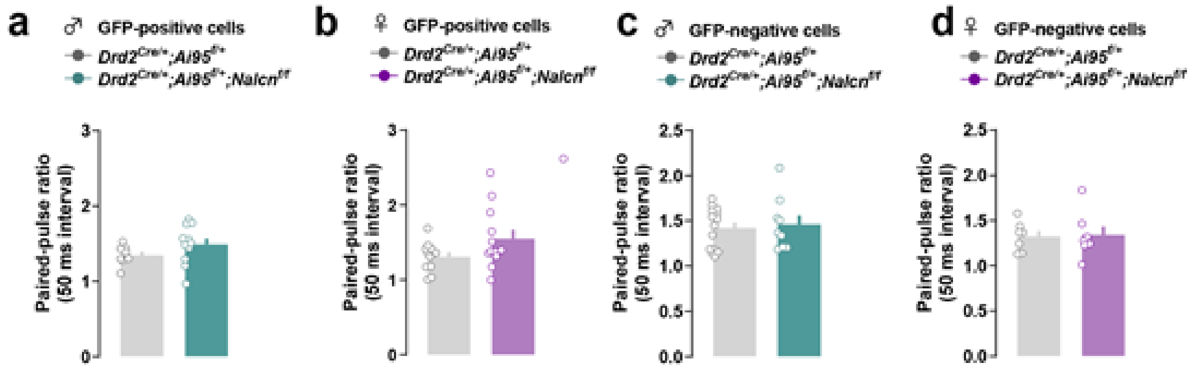

**Supplemental Figure 8: Paired-pulse ratio in SPNs of control and cKO mice.** (**a-d**) Paired-pulse ratio measured in *Drd2*-positive SPNs of male (**a**) and female (**b**) and in *Drd2*-negative SPNs of male (**c**) and female (**d**) mice. Values are represented as mean ± SEM. (detailed statistical analysis in Supplemental Table 2: S8a-d).

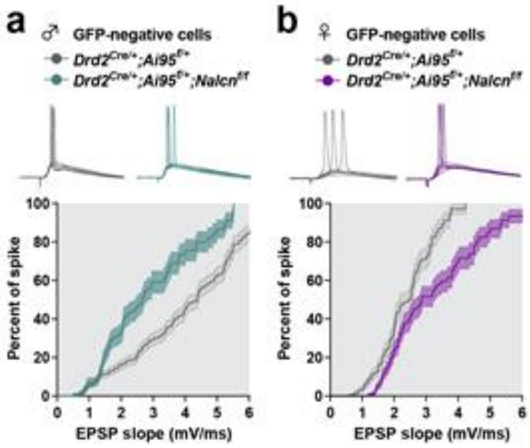

**Supplemental Figure 9: Excitability of *Drd2*-negative SPNs in control and cKO mice.** (**a**-**b**) Survival curves of spike occurrence as a function of EPSP slope (mV/ms) for *Drd2*-negative SPNs in male (**a**) and female (**b**) mice. Upper insets: illustrative traces of neurons clamped at -60 mV with voltage responses for electrical stimulations of excitatory inputs with various intensities evoking EPSPs with or without spike. Values are represented as survival curves with mean ± SEM. (detailed statistical analysis in Supplemental Table 2: S9a-b).

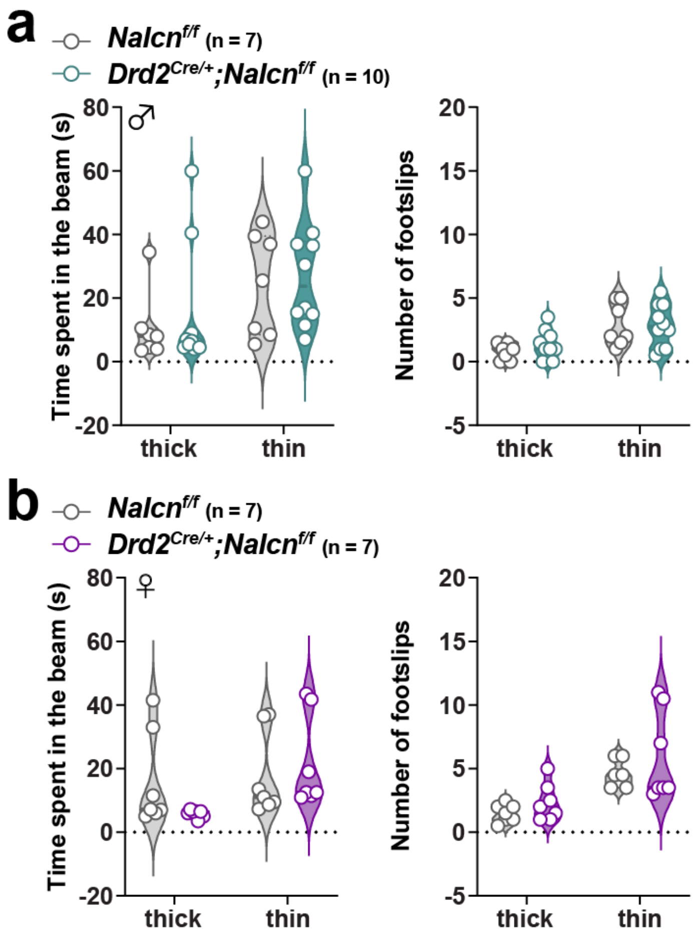

**Supplemental Figure 10: Analysis of motor balance and coordination in control and cKO mice.** (**a-b**) Time spent and number of foot slips in both thick and thin beams in male (**a**) and female (**b**) in *Nalcn*$^{f/f}$ (control) and *Drd2*$^{Cre/+}$;*Nalcn*$^{f/f}$ (cKO) mice (detailed statistical analysis in Supplemental Table 2: S10a-b).

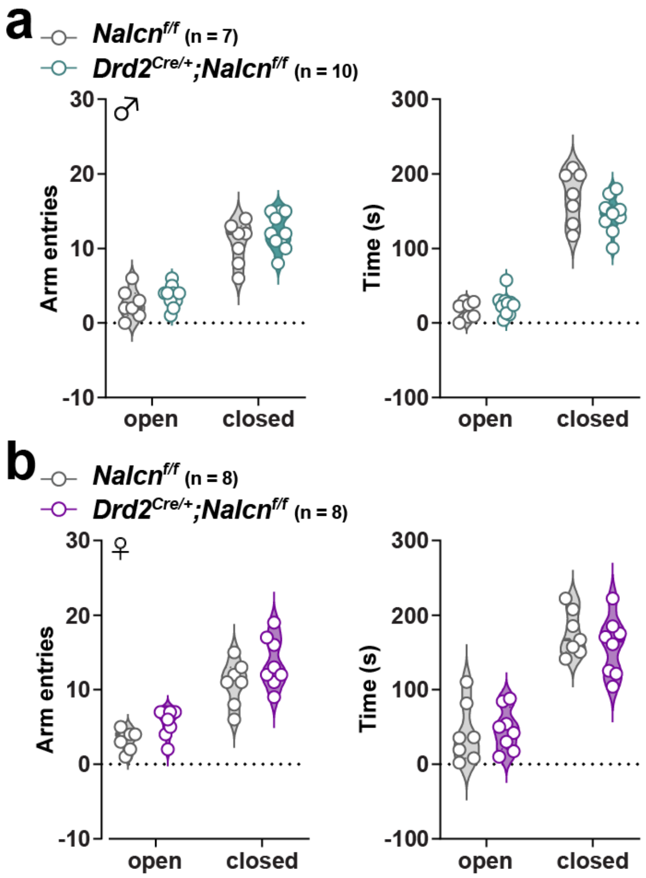

**Supplemental Figure 11: Anxiety-like behavior in control and cKO mice.** Number of entries and time spent on each arm during 5 minutes in the elevated plus maze in male (**a**) and female (**b**) *Nalcn*$^{f/f}$ (control) and *Drd2*$^{Cre/+}$;*Nalcn*$^{f/f}$ (cKO) mice (detailed statistical analysis in Supplemental Table 2: S11a-b)

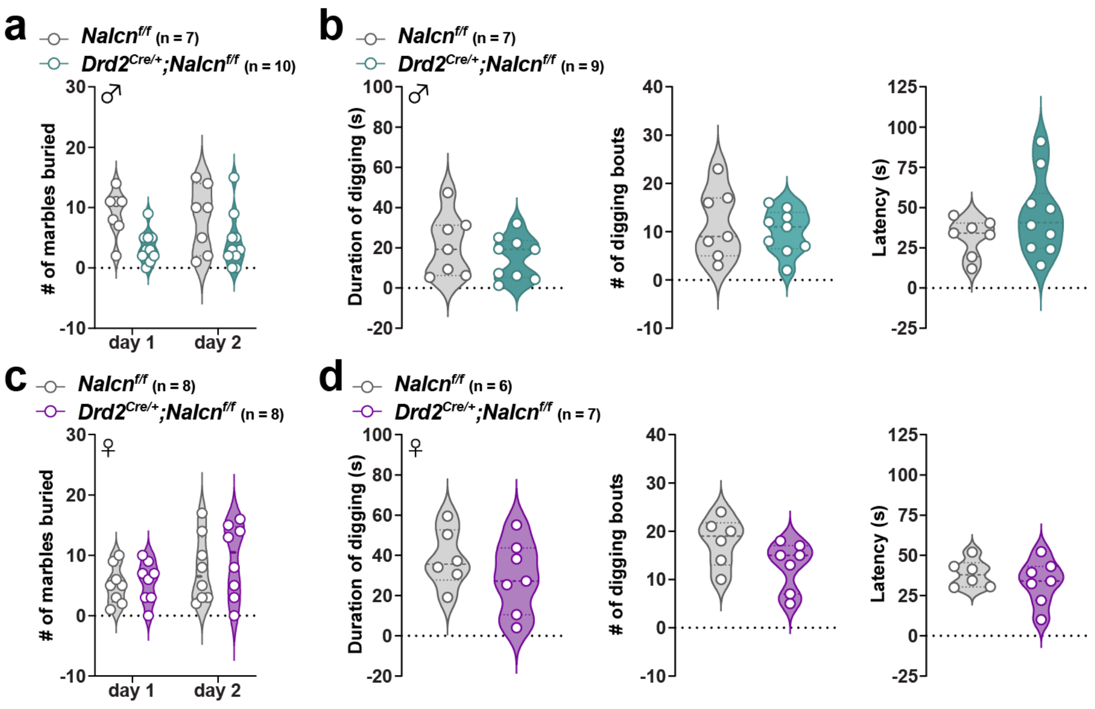

**Supplemental Figure 12: Analysis of digging behaviors in control and cKO mice.** (**a** and **c**) Number of marbles buried after 20 minutes on day 1 and 2 of marble burying test in male (**a**) and female (**c**) *Nalcn*$^{f/f}$ (control) and *Drd2*$^{Cre/+}$;*Nalcn*$^{f/f}$ (cKO) mice. (**b** and **d**) Duration, number of digging bouts and latency to start digging in male (**b**) and female (**d**) *Nalcn*$^{f/f}$ (control) and *Drd2*$^{Cre/+}$;*Nalcn*$^{f/f}$ (cKO) (detailed statistical analysis in Supplemental Table 2: S12a-d).

### Supplementary Table 1: List of Primary Antibodies

| Antigen | Species | Dilution | Supplier/Catalog no./References) |
|---|---|---|---|
| DARPP-32 | Rabbit | 1:1000 (IF); XX (WB) | Cell Signaling Technology (#2306) |
| GFP | Chicken | 1:1000 (IF) | Life Technologies (#A10262) |
| VGAT | Rabbit | 1:1000 (IF) | Synaptic System (#135303) |
| GAD67 | Mouse | 1:1000 (WB) | Millipore (#MAB5406) |
| GAD65 | Rabbit | 1:1000 (WB) | Synaptic System (#198102) |
| STEP | Mouse | 1:1000 (IF) | Cell Signaling Technology (#43965) |
| TH | Rabbit | 1:1000 (IF); 1:5000 (WB) | Millipore (#AB152) |
| DAT | Rat | 1:500 (IF); 1:1000 (WB) | Millipore (#MAB369) |
| VMAT2 | Rabbit | 1:1000 (WB) | Synaptic System (#138302) |
| Parvalbumin | Rabbit | 1:1000 (IF) | Swant (#PV25) |
| NPY | Rabbit | 1:500 (IF) | Abcam (#ab10980) |
| VAChT | Rabbit | 1:500 (IF) | Synaptic System (#139103) |
| β-actin | Mouse | 1:40000 (WB) | Abcam (#AB6276) |
| cFOS | Rat | 1: 2000 (IF) | Synaptic System (#2277) |
| pS6-235/236 | Rabbit | 1:500 (IF) | Cell Signaling Technology (#2211) |
| pH3-S10 | Rabbit | 1:500 (IF) | Millipore (#06-150) |

**Supplementary Table 2: Statistical analysis**

| Figure | Measure | | Groups | Statistical Analysis |
|---|---|---|---|---|
| 2a<br><br><br><br><br>2a inset | Instantaneous current (nA) as function of voltage steps<br><br><br><br>Conductance (mV/pA) | *Drd2*-positive SPNs (male) | $Drd2^{Cre/+}$;$Ai95^{f/+}$ (neurons n = 8)<br>$Drd2^{Cre/+}$;$Ai95^{f/+}$;$Nalcn^{f/f}$ (neurons n = 12) | **Two-way ANOVA repeated measures**<br>Genotype<br>$F_{(1, 18)} = 30.4, p < 0.0001$***<br><br>**Unpaired t-test**<br>$t_{18} = 4.376, p = 0.0024$** |
| 2b<br><br><br><br><br>2b inset | Instantaneous current (nA) as function of voltage steps<br><br><br><br>Conductance (mV/pA) | *Drd2*-positive SPNs (female) | $Drd2^{Cre/+}$;$Ai95^{f/+}$ (neurons n = 11)<br>$Drd2^{Cre/+}$;$Ai95^{f/+}$;$Nalcn^{f/f}$ (neurons n = 12) | **Two-way ANOVA repeated measures**<br>Genotype<br>$F_{(1, 21)} = 3.51, p = 0.075$<br><br>**Unpaired t-test**<br>$T_{21} = 1.813, p = 0.0842$ |
| 3a | RMP (mV) | *Drd2*-positive SPNs (male) | $Drd2^{Cre/+}$;$Ai95^{f/+}$ (neurons n = 9)<br>$Drd2^{Cre/+}$;$Ai95^{f/+}$;$Nalcn^{f/f}$ (neurons n = 11) | **Unpaired t-test**<br>$t_{18} = 2.121, p = 0.0481$* |
| 3b | I/V curve | | | **Two-way ANOVA repeated measures**<br>Genotype<br>$F_{(1, 18)} = 7.617, p = 0.0129$* |
| 3c | Resistance (mOhm) | | | **Unpaired t-test**<br>$t_{18} = 2.515, p = 0.0216$* |
| 3d | I/F spike curve | | | **Two-way ANOVA repeated measures**<br>Genotype<br>$F_{(1, 18)} = 0.280, p = 0.6029$ |
| 3e | Rheobase (pA) | | | **Unpaired t-test**<br>$t_{18} = 0.6399, p = 0.5303$ |
| 3f | RMP (mV) | *Drd2*-positive SPNs (female) | $Drd2^{Cre/+}$;$Ai95^{f/+}$ (neurons n = 11)<br>$Drd2^{Cre/+}$;$Ai95^{f/+}$;$Nalcn^{f/f}$ (neurons n = 9) | **Mann-Whitney test**<br>$U = 23, p = 0.0481$* |
| 3g | I/V curve | | | **Two-way ANOVA repeated measures**<br>Genotype<br>$F_{(1, 18)} = 1.937, p = 0.1809$ |
| 3h | Resistance (mOhm) | | | **Unpaired t-test**<br>$t_{18} = 1.898, p = 0.0739$ |
| 3i | I/F spike curve | | | **Two-way ANOVA repeated measures**<br>Genotype<br>$F_{(1, 18)} = 3.962, p = 0.0619$ |

| | | | | |
|---|---|---|---|---|
| 3j | Rheobase (pA) | | | **Unpaired t-test**<br>$t_{18}$ = 2.169, p = 0.0437* |
| $4a_1$ | sEPSCs inter-event interval (ms) distribution | *Drd2*-positive SPNs (male) | *Drd2$^{Cre/+}$;Ai95$^{f/+}$*<br>(neurons n = 9)<br>*Drd2$^{Cre/+}$;Ai95$^{f/+}$;Nalcn$^{f/f}$*<br>(neurons n = 14) | **Two-way ANOVA repeated measures**<br>Genotype<br>$F_{(1, 23)}$ = 9.907, p = 0.0017** |
| $4a_1$ inset | mean sEPSC events / s | | | **Mann-Whitney test**<br>U = 53.5, p = 0.5676 |
| $4a_2$ | sEPSCs amplitude (pA) distribution | | | **Two-way ANOVA repeated measures**<br>Genotype<br>$F_{(1, 23)}$ = 62.91, p < 0.0001*** |
| $4a_2$ inset | sEPSCs amplitude (pA) | | | **Unpaired t-test**<br>$t_{21}$ = 2.772, p = 0.0114* |
| $4a_3$ | Rise time (ms)<br>Decay time (ms) | | | **Unpaired t-test**<br>$t_{21}$ = 3.609, p = 0.0016**<br>$t_{21}$ = 0.4837, p = 0.6336 |
| $4b_1$ | sEPSCs inter-event interval (ms) distribution | *Drd2*-positive SPNs (female) | *Drd2$^{Cre/+}$;Ai95$^{f/+}$*<br>(neurons n = 9)<br>*Drd2$^{Cre/+}$;Ai95$^{f/+}$;Nalcn$^{f/f}$*<br>(neurons n = 12) | **Two-way ANOVA repeated measures**<br>Genotype<br>$F_{(1, 19)}$ = 18.57, p < 0.0001*** |
| $4b_1$ inset | mean sEPSC events / s | | | **Mann-Whitney test**<br>U = 36, p = 0.2121 |
| $4b_2$ | sEPSCs amplitude (pA) distribution | | | **Two-way ANOVA repeated measures**<br>Genotype<br>$F_{(1, 19)}$ = 3.892, p = 0.0491* |
| $4b_2$ inset | sEPSCs amplitude (pA) | | | **Unpaired t-test**<br>$t_{19}$ = 0.5956, p = 0.5585 |
| $4b_3$ | Rise time (ms)<br>Decay time (ms) | | | **Unpaired t-test**<br>$t_{19}$ = 0.6883, p = 0.4996<br>$t_{19}$ = 3.472, p = 0.0026** |
| 5a | EPSP/spike coupling | *Drd2*-positive SPNs (male) | *Drd2$^{Cre/+}$;Ai95$^{f/+}$*<br>(neurons n = 8)<br>*Drd2$^{Cre/+}$;Ai95$^{f/+}$;Nalcn$^{f/f}$*<br>(neurons n = 12) | **Log-rank (Mantel-Cox) test**<br>x2 = 4.59, p = 0.0322* |
| 5b | EPSP/spike coupling | *Drd2*-positive SPNs (female) | *Drd2$^{Cre/+}$;Ai95$^{f/+}$*<br>(neurons n = 10)<br>*Drd2$^{Cre/+}$;Ai95$^{f/+}$;Nalcn$^{f/f}$*<br>(neurons n = 9) | **Log-rank (Mantel-Cox) test**<br>x2 = 17.97, p < 0.0001*** |
| 6a | GFP/cFOS | male | *Drd2$^{Cre/+}$;Ai95$^{f/+}$*<br>(saline: mice n = 4)<br>*Drd2$^{Cre/+}$;Ai95$^{f/+}$;Nalcn$^{f/f}$*<br>(saline: mice = 3) | **Two-way ANOVA**<br>Genotype<br>$F_{(1, 11)}$ = 0.9764, p = 0.3443 |

| | | | | |
|---|---|---|---|---|
| | | | *Drd2*$^{Cre/+}$;*Ai95*$^{f/+}$ (eticlopride: mice n = 5) *Drd2*$^{Cre/+}$;*Ai95*$^{f/+}$;*Nalcn*$^{f/f}$ (eticlopride: mice n = 3) | |
| | GFP/pS6-S235/236 | | | $F_{(1, 11)}$ = 2.562, p = 0.1377 |
| | GFP/cFOS/pS6-S235/236t | | | $F_{(1, 11)}$ = 1.407, p = 0.2605 |
| 6b | GFP/cFOS/pH3-S10 | male | *Drd2*$^{Cre/+}$;*Ai95*$^{f/+}$ (saline: mice n = 4) *Drd2*$^{Cre/+}$;*Ai95*$^{f/+}$;*Nalcn*$^{f/f}$ (saline: mice = 3) *Drd2*$^{Cre/+}$;*Ai95*$^{f/+}$ (eticlopride: mice n = 5) *Drd2*$^{Cre/+}$;*Ai95*$^{f/+}$;*Nalcn*$^{f/f}$ (eticlopride: mice n = 3) | **Two-way ANOVA** Genotype $F_{(1, 11)}$ = 0.9432, p = 0.3523 |
| 6c | GFP/cFOS | female | *Drd2*$^{Cre/+}$;*Ai95*$^{f/+}$ (saline: mice n = 3) *Drd2*$^{Cre/+}$;*Ai95*$^{f/+}$;*Nalcn*$^{f/f}$ (saline: mice = 4) *Drd2*$^{Cre/+}$;*Ai95*$^{f/+}$ (eticlopride: mice n = 8) *Drd2*$^{Cre/+}$;*Ai95*$^{f/+}$;*Nalcn*$^{f/f}$ (eticlopride: mice n = 7) | **Two-way ANOVA** Genotype $F_{(1, 18)}$ = 1.759, p = 0.2013 |
| | GFP/pS6-S235/236 | | | $F_{(1, 18)}$ = 6.843, p = 0.0175* |
| | GFP/cFOS/pS6-S235/236t | | | $F_{(1, 18)}$ = 5.115, p = 0.0371* |
| 6d | GFP/cFOS/pH3-S10 | female | *Drd2*$^{Cre/+}$;*Ai95*$^{f/+}$ (saline: mice n = 3) *Drd2*$^{Cre/+}$;*Ai95*$^{f/+}$;*Nalcn*$^{f/f}$ (saline: mice = 4) *Drd2*$^{Cre/+}$;*Ai95*$^{f/+}$ (eticlopride: mice n = 8) *Drd2*$^{Cre/+}$;*Ai95*$^{f/+}$;*Nalcn*$^{f/f}$ (eticlopride: mice n = 7) | **Two-way ANOVA** Genotype $F_{(1, 18)}$ = 4.967, p = 0.0396* |
| 7a | Locomotion | male | *Nalcn*$^{f/f}$ (mice n = 7) *Drd2*$^{Cre/+}$;*Nalcn*$^{f/f}$ (mice n = 10) | **Two-way ANOVA repeated measures** Genotype $F_{(1, 15)}$ = 12.39 , p = 0.0031** |
| | Total | | | **Unpaired t-test** $t_{15}$ = 3.521, p = 0.0031** |
| 7c | Rearings | male | *Nalcn*$^{f/f}$ (mice n = 7) *Drd2*$^{Cre/+}$;*Nalcn*$^{f/f}$ (mice n = 10) | **Two-way ANOVA repeated measures** Genotype $F_{(1, 15)}$ = 5.756 , p = 0.0299* |
| | Total | | | **Unpaired t-test** $t_{15}$ = 2.399, p = 0.0299* |
| 7b | Locomotion | female | *Nalcn*$^{f/f}$ (mice n = 8) | **Two-way ANOVA repeated measures** |

| | | | | |
|---|---|---|---|---|
| | Total | | *Drd2*$^{Cre/+}$;*Nalcn*$^{f/f}$<br>(mice n = 8) | Genotype<br>$F_{(1, 14)}$ = 1.676 , p = 0.2163<br><br>**Unpaired t-test**<br>$t_{14}$ = 1.295, p = 0.2163 |
| 7d | Rearings<br><br><br><br><br>Total | female | *Nalcn*$^{f/f}$<br>(mice n = 8)<br>*Drd2*$^{Cre/+}$;*Nalcn*$^{f/f}$<br>(mice n = 8) | **Two-way ANOVA repeated measures**<br>Genotype<br>$F_{(1, 14)}$ = 0.2211 , p = 0.6455<br><br>**Unpaired t-test**<br>$t_{14}$ = 0.4702, p = 0.6455 |
| 7e | Total lever presses | male | *Nalcn*$^{f/f}$<br>(mice n = 6)<br>*Drd2*$^{Cre/+}$;*Nalcn*$^{f/f}$<br>(mice n = 12) | **FR1: Three-way ANOVA repeated measures**<br>Genotype<br>$F_{(1, 30)}$ = 0.8791 , p = 0.3559<br><br>**FR5: Three-way ANOVA repeated measures**<br>Genotype<br>$F_{(1, 32)}$ = 8.039 , p = 0.0079** |
| 7f | Pellet consumed | male | *Nalcn*$^{f/f}$<br>(mice n = 6)<br>*Drd2*$^{Cre/+}$;*Nalcn*$^{f/f}$<br>(mice n = 12) | **FR1: Two-way ANOVA repeated measures**<br>Genotype<br>$F_{(1, 16)}$ = 3.786 , p = 0.0695<br><br>**FR5: Two-way ANOVA repeated measures**<br>Genotype<br>$F_{(1, 16)}$ = 16.23 , p = 0.001*** |
| 7g | Cumulative lever press during time out | male | *Nalcn*$^{f/f}$<br>(mice n = 6)<br>*Drd2*$^{Cre/+}$;*Nalcn*$^{f/f}$<br>(mice n = 12) | **FR1: Three-way ANOVA repeated measures**<br>Genotype<br>$F_{(1, 32)}$ = 0.2660 , p = 0.6096<br><br>**FR5: Three-way ANOVA repeated measures**<br>Genotype<br>$F_{(1, 32)}$ = 5.048 , p = 0.0317* |
| 7h | Breaking point | male | *Nalcn*$^{f/f}$<br>(mice n = 6)<br>*Drd2*$^{Cre/+}$;*Nalcn*$^{f/f}$<br>(mice n = 12) | **Unpaired t-test**<br>$t_{16}$ = 0.9563, p = 0.3531 |
| 7i | Total lever presses | female | *Nalcn*$^{f/f}$<br>(mice n = 6)<br>*Drd2*$^{Cre/+}$;*Nalcn*$^{f/f}$<br>(mice n = 4) | **FR1: Three-way ANOVA repeated measures**<br>Genotype<br>$F_{(1, 16)}$ = 0.9388 , p = 0.347<br><br>**FR5: Three-way ANOVA repeated measures**<br>Genotype |

| | | | | |
|---|---|---|---|---|
| | | | | $F_{(1, 16)} = 0.0276$ , $p = 0.8701$ |
| 7j | Pellet consumed | female | *Nalcn*$^{f/f}$<br>(mice n = 6)<br>*Drd2*$^{Cre/+}$;*Nalcn*$^{f/f}$<br>(mice n = 4) | **FR1: Two-way ANOVA repeated measures**<br>Genotype<br>$F_{(1, 8)} = 0.186$ , $p = 0.6777$<br><br>**FR5: Two-way ANOVA repeated measures**<br>Genotype<br>$F_{(1, 8)} = 0.7482$ , $p = 0.4122$ |
| 7k | Cumulative lever press during time out | female | *Nalcn*$^{f/f}$<br>(mice n = 6)<br>*Drd2*$^{Cre/+}$;*Nalcn*$^{f/f}$<br>(mice n = 4) | **FR1: Three-way ANOVA repeated measures**<br>Genotype<br>$F_{(1, 16)} = 0.6256$ , $p = 0.4405$<br><br>**FR5: Three-way ANOVA repeated measures**<br>Genotype<br>$F_{(1, 32)} = 0.1414$ , $p = 0.7118$ |
| 7l | Breaking point | female | *Nalcn*$^{f/f}$<br>(mice n = 6)<br>*Drd2*$^{Cre/+}$;*Nalcn*$^{f/f}$<br>(mice n = 4) | **Unpaired t-test**<br>$t_8 = 0.01628$, $p = 0.9874$ |
| 8b | Avoidance probability | male | *Nalcn*$^{f/f}$<br>(mice n = 11)<br>*Drd2*$^{Cre/+}$;*Nalcn*$^{f/f}$<br>(mice n = 8) | **CS+: Two-way ANOVA repeated measures**<br>Genotype<br>$F_{(1, 17)} = 1.953$ , $p = 0.1802$ |
| 8c | Defensive probability | male | *Nalcn*$^{f/f}$<br>(mice n = 11)<br>*Drd2*$^{Cre/+}$;*Nalcn*$^{f/f}$<br>(mice n = 8) | **CS-: Two-way ANOVA repeated measures**<br>Genotype<br>$F_{(1, 17)} = 25.96$ , $p < 0.001$*** |
| 8d | Discrimination performance | male | *Nalcn*$^{f/f}$<br>(mice n = 11)<br>*Drd2*$^{Cre/+}$;*Nalcn*$^{f/f}$<br>(mice n = 8) | **Two-way ANOVA repeated measures**<br>Genotype<br>$F_{(1, 17)} = 13.35$ , $p = 0.002$** |
| 8e | Freezing probability | male | *Nalcn*$^{f/f}$<br>(mice n = 10)<br>*Drd2*$^{Cre/+}$;*Nalcn*$^{f/f}$<br>(mice n = 12) | **Two-way ANOVA repeated measures**<br>Genotype<br>$F_{(1, 20)} = 1.935$ , $p = 0.1795$ |
| S1c | DARPP-32<br><br><br><br><br>GAD67 | | *Nalcn*$^{f/f}$<br>(mice n = 6)<br>*Drd2*$^{Cre/+}$;*Nalcn*$^{f/f}$<br>(mice n = 6)<br><br>*Nalcn*$^{f/f}$ | **Unpaired t-test**<br>$t_{10} = 0.5534$, $p = 0.5921$<br><br><br><br>$t_9 = 0.0974$, $p = 0.9245$ |

| | | | | |
|---|---|---|---|---|
| | GAD65 | | (mice n = 6)<br>*Drd2*$^{Cre/+}$;*Nalcn*$^{f/f}$<br>(mice n = 5)<br><br>*Nalcn*$^{f/f}$<br>(mice n = 7)<br>*Drd2*$^{Cre/+}$;*Nalcn*$^{f/f}$<br>(mice n = 7) | $t_{12}$ = 0.9151, p = 0.3782 |
| S2 | TH<br><br><br><br><br>VMAT2<br><br><br><br><br>DAT | | *Nalcn*$^{f/f}$<br>(mice n = 6)<br>*Drd2*$^{Cre/+}$;*Nalcn*$^{f/f}$<br>(mice n = 6)<br><br>*Nalcn*$^{f/f}$<br>(mice n = 6)<br>*Drd2*$^{Cre/+}$;*Nalcn*$^{f/f}$<br>(mice n = 6)<br><br>*Nalcn*$^{f/f}$<br>(mice n = 5)<br>*Drd2*$^{Cre/+}$;*Nalcn*$^{f/f}$<br>(mice n = 6) | **Unpaired t-test**<br>$t_{10}$ = 0.4014, p = 0.6966<br><br><br><br>$t_{10}$ = 0.8786, p = 0.4002<br><br><br><br><br>$t_9$ = 0.6968, p = 0.5035 |
| S4a | Instantaneous current (nA) as function of voltage steps | *Drd2*-positive SPNs (male) | *Drd2*$^{Cre/+}$;*Ai95*$^{f/+}$<br>(neurons n = 8)<br>*Drd2*$^{Cre/+}$;*Ai95*$^{f/+}$;*Nalcn*$^{f/f}$<br>(neurons n = 12) | **Two-way ANOVA repeated measures**<br>Genotype<br>$F_{(1, 18)}$ = 1.2848, p = 0.2719 |
| S4b | Instantaneous current (nA) as function of voltage steps | *Drd2*-positive SPNs (female) | *Drd2*$^{Cre/+}$;*Ai95*$^{f/+}$<br>(neurons n = 11)<br>*Drd2*$^{Cre/+}$;*Ai95*$^{f/+}$;*Nalcn*$^{f/f}$<br>(neurons n = 12) | **Two-way ANOVA repeated measures**<br>Genotype<br>$F_{(1, 21)}$ = 0.359, p = 0.5553 |
| S4c | Instantaneous current (nA) as function of voltage steps | *Drd2*-negative SPNs (male) | *Drd2*$^{Cre/+}$;*Ai95*$^{f/+}$<br>(neurons n = 13)<br>*Drd2*$^{Cre/+}$;*Ai95*$^{f/+}$;*Nalcn*$^{f/f}$<br>(neurons n = 11) | **Two-way ANOVA repeated measures**<br>Genotype<br>$F_{(1, 22)}$ = 3.5, p = 0.0747 |
| S4d | Instantaneous current (nA) as function of voltage steps | *Drd2*-negative SPNs (female) | *Drd2*$^{Cre/+}$;*Ai95*$^{f/+}$<br>(neurons n = 8)<br>*Drd2*$^{Cre/+}$;*Ai95*$^{f/+}$;*Nalcn*$^{f/f}$<br>(neurons n = 8) | **Two-way ANOVA repeated measures**<br>Genotype<br>$F_{(1, 14)}$ = 6.378, p = 0.0242* |
| S5a<br><br><br><br><br>inset | Instantaneous current (A) as function of voltage steps<br><br><br>Conductance (mV/pA) | *Drd2*-negative SPNs (male) | *Drd2*$^{Cre/+}$;*Ai95*$^{f/+}$<br>(neurons n = 13)<br>*Drd2*$^{Cre/+}$;*Ai95*$^{f/+}$;*Nalcn*$^{f/f}$<br>(neurons n = 11) | **Two-way ANOVA repeated measures**<br>Genotype<br>$F_{(1, 22)}$ = 2.803, p =0.1082<br><br>**Unpaired t-test**<br>$T_{22}$ = 1.237, p = 0.2289 |
| S5b | Instantaneous current (A) as function of voltage steps | *Drd2*-negative SPNs (female) | *Drd2*$^{Cre/+}$;*Ai95*$^{f/+}$<br>(neurons n = 8)<br>*Drd2*$^{Cre/+}$;*Ai95*$^{f/+}$;*Nalcn*$^{f/f}$<br>(neurons n = 8) | **Two-way ANOVA repeated measures**<br>Genotype<br>$F_{(1, 14)}$ = 9.2e-005, p = 0.9925 |

| | | | | |
|---|---|---|---|---|
| inset | Conductance (mV/pA) | | | **Unpaired t-test**<br>$t_{14} = 0.3225$, $p = 0.7518$ |
| S6a | RMP (mV) | *Drd2*-negative SPNs (male) | *Drd2*$^{Cre/+}$;*Ai95*$^{f/+}$ (neurons n = 13)<br>*Drd2*$^{Cre/+}$;*Ai95*$^{f/+}$;*Nalcn*$^{f/f}$ (neurons n = 9) | **Unpaired t-test**<br>$t_{20} = 0.8536$, $p = 0.4035$ |
| S6b | I/V curve | | | **Two-way ANOVA repeated measures**<br>Genotype<br>$F_{(1, 20)} = 0.021$, $p = 0.8860$ |
| S6c | Resistance (mOhm) | | | **Unpaired t-test**<br>$t_{20} = 0.1865$, $p = 0.8540$ |
| S6d | I/F spike curve | | | **Two-way ANOVA repeated measures**<br>Genotype<br>$F_{(1, 20)} = 3.881$, $p = 0.0628$ |
| S6e | Rheobase (pA) | | | **Unpaired t-test**<br>$t_{20} = 0.673$, $p = 0.5087$ |
| S6f | RMP (mV) | *Drd2*-negative SPNs (female) | *Drd2*$^{Cre/+}$;*Ai95*$^{f/+}$ (neurons n = 8)<br>*Drd2*$^{Cre/+}$;*Ai95*$^{f/+}$;*Nalcn*$^{f/f}$ (neurons n = 9) | **Unpaired t-test**<br>$t_{15} = 0.9442$, $p = 0.36$ |
| S6g | I/V curve | | | **Two-way ANOVA repeated measures**<br>Genotype<br>$F_{(1, 15)} = 0.333$, $p = 0.5724$ |
| S6h | Resistance (mOhm) | | | **Unpaired t-test**<br>$t_{15} = 0.7836$, $p = 0.4455$ |
| S6i | I/F spike curve | | | **Two-way ANOVA repeated measures**<br>Genotype<br>$F_{(1, 15)} = 0.642$, $p = 0.4353$ |
| S6j | Rheobase (pA) | | | **Unpaired t-test**<br>$t_{15} = 1.628$, $p = 0.1244$ |
| S7a$_1$<br><br>inset<br><br>S7a$_2$<br><br>inset | sEPSCs inter-event interval (ms) distribution<br><br>mean sEPSC events / s<br><br>sEPSCs amplitude (pA) distribution<br><br>sEPSCs amplitude (pA) | *Drd2*-negative SPNs (male) | *Drd2*$^{Cre/+}$;*Ai95*$^{f/+}$ (neurons n = 14)<br>*Drd2*$^{Cre/+}$;*Ai95*$^{f/+}$;*Nalcn*$^{f/f}$ (neurons n = 9) | **Two-way ANOVA repeated measures**<br>Genotype<br>$F_{(1, 21)} = 33.13$, $p < 0.0001$***<br><br>**Unpaired t-test**<br>$t_{21} = 1.007$, $p = 0.3254$<br><br>**Two-way ANOVA repeated measures**<br>Genotype<br>$F_{(1, 23)} = 0.4012$, $p = 0.5267$<br><br>**Unpaired t-test**<br>$t_{21} = 0.1536$, $p = 0.8794$ |

| | | | | |
|---|---|---|---|---|
| S7a$_3$ | Rise time (ms)<br>Decay time (ms) | | | **Unpaired t-test**<br>$t_{21}$ = 0.03681, p = 0.971<br>$t_{21}$ = 0.2335, p = 0.8176 |
| S7b$_1$<br><br><br><br><br>inset<br><br><br>S7b$_2$<br><br><br><br><br>inset<br><br><br>S7b$_3$ | sEPSCs inter-event interval (ms) distribution<br><br><br><br><br>mean sEPSC events / s<br><br><br>sEPSCs amplitude (pA) distribution<br><br><br><br>inset<br>sEPSCs amplitude (pA)<br><br><br>Rise time (ms)<br>Decay time (ms) | *Drd2*-negative SPNs (female) | *Drd2*$^{Cre/+}$;*Ai95*$^{f/+}$<br>(neurons n = 8)<br>*Drd2*$^{Cre/+}$;*Ai95*$^{f/+}$;*Nalcn*$^{f/f}$<br>(neurons n = 8) | **Two-way ANOVA repeated measures**<br>Genotype<br>$F_{(1, 14)}$ = 1.561, p = 0.2119<br><br>**Mann-Whitney test**<br>$t_{14}$ = 0.1.226, p = 0.2404<br><br>**Two-way ANOVA repeated measures**<br>Genotype<br>$F_{(1, 14)}$ = 1.303, p = 0.2545<br><br>**Unpaired t-test**<br>$t_{14}$ = 0.3521, p = 0.73<br><br>**Unpaired t-test**<br>$t_{14}$ = 0.293, p = 0.7738<br>$t_{14}$ = 0.7688, p = 0.4548 |
| S8a | Paired-pulse ratio | *Drd2*-positive SPNs (male) | *Drd2*$^{Cre/+}$;*Ai95*$^{f/+}$<br>(neurons n = 8)<br>*Drd2*$^{Cre/+}$;*Ai95*$^{f/+}$;*Nalcn*$^{f/f}$<br>(neurons n = 11) | **Unpaired t-test**<br>$t_{17}$ = 1.442, p = 0.1674 |
| S8b | Paired-pulse ratio | *Drd2*-positive SPNs (female) | *Drd2*$^{Cre/+}$;*Ai95*$^{f/+}$<br>(neurons n = 11)<br>*Drd2*$^{Cre/+}$;*Ai95*$^{f/+}$;*Nalcn*$^{f/f}$<br>(neurons n = 12) | **Unpaired t-test**<br>$t_{21}$ = 1.817, p = 0.0877 |
| S8c | Paired-pulse ratio | *Drd2*-negative SPNs (male) | *Drd2*$^{Cre/+}$;*Ai95*$^{f/+}$<br>(neurons n = 14)<br>*Drd2*$^{Cre/+}$;*Ai95*$^{f/+}$;*Nalcn*$^{f/f}$<br>(neurons n = 9) | **Unpaired t-test**<br>$t_{21}$ = 0.3917, p = 0.6992 |
| S8d | Paired-pulse ratio | *Drd2*-negative SPNs (female) | *Drd2*$^{Cre/+}$;*Ai95*$^{f/+}$<br>(neurons n = 7)<br>*Drd2*$^{Cre/+}$;*Ai95*$^{f/+}$;*Nalcn*$^{f/f}$<br>(neurons n = 7) | **Unpaired t-test**<br>$t_{12}$ = 0.1345, p = 0.8955 |
| S9a | EPSP/spike coupling | *Drd2*-negative SPNs (male) | *Drd2*$^{Cre/+}$;*Ai95*$^{f/+}$<br>(neurons n = 11)<br>*Drd2*$^{Cre/+}$;*Ai95*$^{f/+}$;*Nalcn*$^{f/f}$<br>(neurons n = 5) | **Log-rank (Mantel-Cox) test**<br>$x2$ = 28.04, p < 0.0001*** |
| S9b | EPSP/spike coupling | *Drd2*-negative SPNs (female) | *Drd2*$^{Cre/+}$;*Ai95*$^{f/+}$<br>(neurons n = 7)<br>*Drd2*$^{Cre/+}$;*Ai95*$^{f/+}$;*Nalcn*$^{f/f}$<br>(neurons n = 7) | **Log-rank (Mantel-Cox) test**<br>$x2$ = 20.14, p < 0.0001*** |
| S10a | Time spent in beam | male | *Nalcn*$^{f/f}$<br>(mice n = 7)<br>*Drd2*$^{Cre/+}$;*Nalcn*$^{f/f}$<br>(mice n = 10) | **Two-way ANOVA repeated measures**<br>Genotype<br>$F_{(1, 15)}$ = 0.2341 , p = 0.6355 |

| | | | | |
|---|---|---|---|---|
| | Number of footslips | | | $F_{(1, 15)}$ = 0.2134 , p = 0.6508 |
| S10b | Time spent in beam<br><br><br><br><br>Number of footslips | female | *Nalcn*$^{f/f}$<br>(mice n = 7)<br>*Drd2*$^{Cre/+}$;*Nalcn*$^{f/f}$<br>(mice n = 7) | **Two-way ANOVA repeated measures**<br>Genotype<br>$F_{(1, 12)}$ = 0.3136 , p = 0.5858<br><br>$F_{(1, 12)}$ = 1.674 , p = 0.2201 |
| S11a | Arm entries<br><br><br><br>Time in arms | male | *Nalcn*$^{f/f}$<br>(mice n = 7)<br>*Drd2*$^{Cre/+}$;*Nalcn*$^{f/f}$<br>(mice n = 10) | **Two-way ANOVA repeated measures**<br>Genotype<br>$F_{(1, 15)}$ = 1.933 , p = 0.1847<br>$F_{(1, 15)}$ = 1.977 , p = 0.9576 |
| S11b | Arm entries<br><br><br><br>Time in arms | female | *Nalcn*$^{f/f}$<br>(mice n = 7)<br>*Drd2*$^{Cre/+}$;*Nalcn*$^{f/f}$<br>(mice n = 8) | **Two-way ANOVA repeated measures**<br>Genotype<br>$F_{(1, 13)}$ = 9.149 , p = 0.0098**<br>$F_{(1, 13)}$ = 0.5433 , p = 0.4742 |
| S12a | Number of marbles buried | male | *Nalcn*$^{f/f}$<br>(mice n = 7)<br>*Drd2*$^{Cre/+}$;*Nalcn*$^{f/f}$<br>(mice n = 10) | **Two-way ANOVA repeated measures**<br>Genotype<br>$F_{(1, 15)}$ = 7.660 , p = 0.0144* |
| S12b | Duration of digging<br><br><br><br><br>Number of digging bouts<br><br>Latency to the 1st digging bouts | male | *Nalcn*$^{f/f}$<br>(mice n = 7)<br>*Drd2*$^{Cre/+}$;*Nalcn*$^{f/f}$<br>(mice n = 9) | **Unpaired t-test**<br>$t_{14}$ = 0.8909, p = 0.3880<br><br><br><br>$t_{14}$ = 0.5305, p = 0.6041<br><br>$t_{14}$ = 1.325, p = 0.2049 |
| S12c | Number of marbles buried | female | *Nalcn*$^{f/f}$<br>(mice n = 8)<br>*Drd2*$^{Cre/+}$;*Nalcn*$^{f/f}$<br>(mice n = 8) | **Two-way ANOVA repeated measures**<br>Genotype<br>$F_{(1, 14)}$ = 0.2667 , p = 0.6136 |
| S12d | Duration of digging<br><br><br><br><br>Number of digging bouts<br><br>Latency to the 1st digging bouts | female | *Nalcn*$^{f/f}$<br>(mice n = 6)<br>*Drd2*$^{Cre/+}$;*Nalcn*$^{f/f}$<br>(mice n = 7) | **Unpaired t-test**<br>$t_{11}$ = 1.012, p = 0.3332<br><br><br><br>$t_{11}$ = 1.78, p = 0.1026<br><br>$t_{11}$ = 0.7984, p = 0.4415 |
